# Archaeal tubulin-like proteins CetZ1 and CetZ2 have opposing effects on cell morphology during the growth cycle of *Haloferax volcanii*

**DOI:** 10.1101/2024.10.29.620987

**Authors:** Hannah J. Brown, Iain G. Duggin

**Affiliations:** The Australian Institute for Microbiology and Infection, University of Technology Sydney, NSW, 2007 Australia

**Keywords:** Archaea, Cytoskeleton, Cell shape, Tubulin, CetZ

## Abstract

CetZs are archaeal tubulin superfamily cytoskeletal proteins implicated in the control of cell shape and motility. In the pleomorphic archaeon *Haloferax volcanii*, CetZ1 is required for the transformation of a discoid, or plate-like, cell morphology to the rod shape during early-mid log phase cultures and for the development of swimming motility. In this study, we found that the paralog CetZ2 is not required for rod development or reversion to plates in log phase but was strongly upregulated later in stationary phase, where it promotes the maintenance of plate cell shape. CetZ2 cytoskeletal structures visualized through the production of a functional fluorescent tagged CetZ2 were most dynamic specifically in mid-stationary phase, where they showed directional movement around the cell edge and other complex cytomotive-like behaviours. Furthermore, in mid-stationary phase, the dynamics of CetZ1 and CetZ2 cytoskeletal structures were specifically dependent on the presence of one another and their GTPase activities that control the polymerization-depolymerization cycle. Together, the results suggest that CetZ2 counteracts the CetZ1-based rod development pathway to maintain plate shape, and they imply that additional stationary-phase factors are also involved. CetZ1 and CetZ2 are co-conserved in many haloarchaea, suggesting that their coordinated functions at different stages of the growth cycle are widely utilized to control cell shapeshifting. More generally, our findings show that CetZ paralogues can have distinct and non-redundant functions, indicating that diversification and specialisation within the tubulin superfamily has occurred multiple times in archaeal evolution.

**Significance:** This study describes the first defined function of a CetZ2 protein, which represents a distinct haloarchaeal protein family within the near-universal tubulin superfamily of cytoskeletal proteins. Using the model archaeal organism *Haloferax volcanii*, we found that CetZ2 maintains the plate or disk-like cell morphology specifically in the stationary phase of the growth cycle, by counteracting CetZ1-based rod development. We also showed for the first time that CetZ2 can form dynamic cytoskeletal filaments that show directional movement around the cell in mid-stationary phase. The CetZ1-CetZ2 interplay we detected specifically at this stage may represent a paradigm for understanding the evolution of antagonistic cytoskeletal functions, such as those that led to a reliance on active and inactive subunits during the early evolution of microtubules in eukaryotes.

## Introduction

CetZs are archaea-specific tubulin superfamily proteins that are structural homologues of eukaryotic tubulin which forms microtubules, and FtsZ, the key cell division protein present in both bacteria and archaea (1, 2). One shared characteristic of these tubulin superfamily proteins is that they polymerise in a GTP-dependent manner, forming filaments or large polymers (3–9), which assemble into larger structures that carry out their biological functions. Both FtsZ and tubulin have a wide diversity of functions which are well studied, including in cell division and chromosome segregation (10, 11), intracellular trafficking and cargo transport (12, 13), and cell motility or migration (14, 15).

CetZs are conserved across many diverse archaea. In Halobacteria (Haloarchaea), it is typical that one species would have multiple paralogues of CetZ, and the most highly conserved of these are CetZ1 and CetZ2 (1, 16, 17), both of which form strong and distinct phylogenetic clusters, suggesting that CetZ1 and CetZ2 could have different functions. While both CetZ1 and CetZ2 have been implicated in the control of shape and motility in *Haloferax volcanii,* only CetZ1 currently has a clearly defined biological function. During standard batch culture, *H. volcanii* transitions from plate-shaped cells to rods between the early and late log phases, but once cells enter stationary phase they revert to plates (18). CetZ1 has shown to be necessary for rod development, not only during early-log (1, 18), but also in other conditions where *H. volcanii* becomes rod-shaped, such as in motility conditions (1, 19, 20) or when depleted of trace metals (18).

CetZ1 localizes in patches around the membrane of cells but is somewhat concentrated at mid-cell and the cell poles, where it is hypothesised to form a cap-like structure in motile rods (1, 20–23). CetZ1 is thought to contribute to motility by generating the rod shape that enables directional and streamlined swimming, as well as by promoting the positioning and assembly of archaella and chemosensory arrays (21). CetZ1 shows dynamic localization during growth and rod development, which is linked to its polymerisation and depolymerisation cycle (1, 22). A point mutation in the GTPase active site (CetZ1.E218A), based on the equivalent mutations in FtsZ and tubulin, causes it to form the expected hyper-stable structures in vivo, thus disrupting CetZ1 assembly dynamics that are important to its function (1). Expression of CetZ1.E218A prevents rod development, even in the presents of the endogenous CetZ1, and produces cells with local regions of high envelope curvature, or ‘jagged’ cells; CetZ1.E218A localizes to the regions of high membrane curvature (1, 22).

While CetZ1 has roles in cell shape control and motility, no clear role of other CetZ paralogues of *H. volcanii* have been identified. Although CetZ2 has been generally implicated in the control of cell shape and motility (1, 21), deletion of *cetZ2* results in no cell division, shape, or motility phenotypes during exponential phase or on soft-agar, respectively (1, 21). However, overexpression of CetZ2 has been demonstrated to induce slight hypermotility and promote assembly of archaellum filaments (21). In addition, expression of CetZ2.E212A, the equivalent GTPase active-site mutant of CetZ1.E218A, prevents rod-development and perturbs motility (1, 21). CetZ2.E212A might therefore interfere with CetZ1 function (1), whether through direct interaction with CetZ1 or indirectly through an unknown mechanism. Together with the knowledge that CetZ2 is only present in Halobacteria that also have CetZ1 (17), the current evidence points towards a potential functional interplay between CetZ2 and CetZ1. One possible explanation for the lack of observable effects on cell shape and motility upon deletion of *cetZ2* is that the CetZ2 protein may be inactive or present in very low abundance in the tested conditions (i.e. early/mid-log growth, motility assays). Here, we show that CetZ2 is strongly upregulated in stationary phase where it maintains plate shape via a direct or indirect counteraction of CetZ1-dependent rod development.

## Results

### CetZ2 is strongly upregulated in stationary phase in H. volcanii

CetZ2 function has previously been investigated during exponential phase growth and motility conditions. However, proteomic data (24) suggested it may be upregulated in stationary phase in *H. volcanii* and *N. magadii*. Here, we used western blotting to detect CetZ2 levels in Hv-YPCab (rich medium), Hv-Cab (casamino acids medium), and Hv-Min (minimal medium) between early log phase and late stationary phase of *H. volcanii* (Fig. 1a). CetZ1 was expressed throughout the growth cycle, though decreased in stationary phase (Fig. 1b). In contrast, there was little production of CetZ2 until stationary phase, where protein levels strongly increase (Fig. 1c). Downregulation of CetZ1 in stationary phase was not dependent on the presence of CetZ2 (Fig. S1a), and upregulation of CetZ2 was not dependant on the presence of CetZ1 (Fig. S1b). The ratios between CetZ1 and CetZ2 and the number of molecules per cell were also measured using CetZ1/2 pure protein standards (Fig. 1d, e, respectively). Wildtype cells in mid-log growth were measured to have on average 5735 molecules of CetZ1 and 142 molecules of CetZ2, giving a ratio of ~41:1 (CetZ1:CetZ2, Fig. 1f). In stationary phase, the amounts averaged 2234 molecules of CetZ1 and 425 molecules of CetZ2 per cell, a ~5:1 ratio (Fig. 1f).

**Figure 1.**
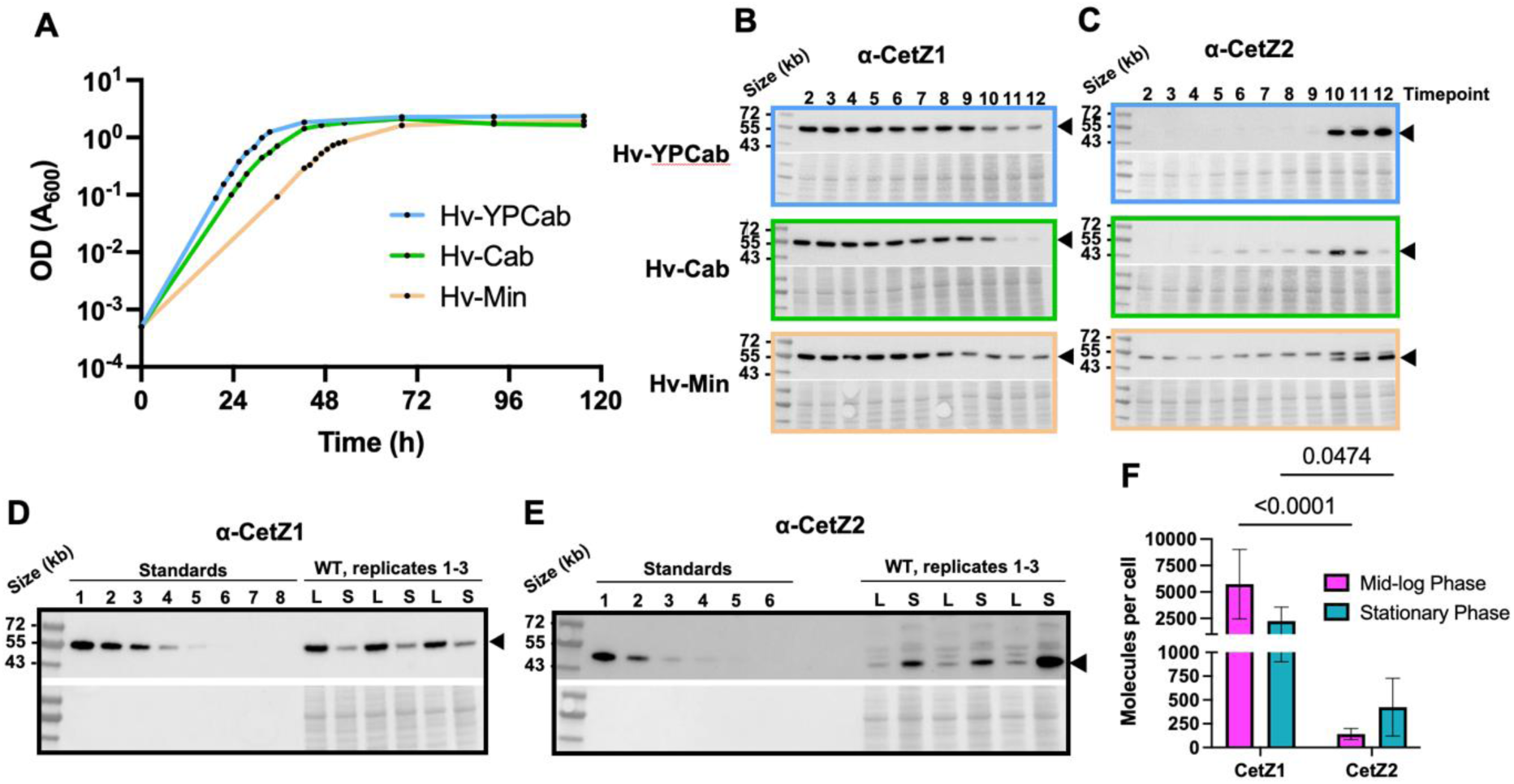
Quantification of CetZ1 and CetZ2 in H. volcanii throughout growth. **A)** Wild-type cells (H26 + pTA962) were grown in Hv-YPCab, Hv-Cab, or Hv-Min from a theoretical starting OD_600_~0.0005 for 120 h. Samples of each culture were taken for western blotting at 12 points along each growth curve, indicated by black data points. **B)** Western blot analysis of wild-type cells grown in **A** using antibodies against CetZ1, and **C)** CetZ2. Samples from timepoints 2-12 of each medium were used for analysis. Ponceau S staining (bottom panels) was carried out to ensure equal sample loading. Solid arrow heads indicate CetZ1/2 bands. **D)** Purified CetZ1 was used to make eight standards of known concentrations. Standard 1 is 50 ng of pure CetZ1, and standards 2-8 are 1:2 serial dilutions of standard 1. Standards were compared to whole-cell lysate samples of wild-type cells to estimate the number of CetZ1 molecules per cell in mid-log (L) and stationary (S) phases using western blot detection of CetZ1. **E)** As in D, except for CetZ2. Standard 1 is 1.25 ng of pure CetZ2, and standards 2-6 are 1:2 serial dilutions of standard 1. In **D** and **E**, six culture replicates were carried out, and the same samples were used for analysis of CetZ1 and CetZ2 expression levels. Solid arrow heads indicate CetZ1/2 bands. **F)** From the results of **D** and **E**, the number of CetZ1/CetZ2 molecules per cell in mid-log and stationary phase were calculated as described in the methods. Mean and standard deviation are shown. Unpaired one-way ANOVA was used as a statistical test.

### CetZ2 contributes to maintenance of plate-shape in stationary phase

CetZ2 has previously been implicated in cell shape through the expression of a GTPase active-site mutant (CetZ2.E212A) (1). This type of T7-loop mutation is expected to inhibit GTPase activity in tubulin-superfamily proteins, producing inactive hyper-stable filaments. Production of CetZ2.E212A resulted in a dominant inhibitory effect, causing distorted cell shapes, similar to the effects of the equivalent mutant CetZ1.E218A, but did not reveal the normal role of CetZ2 (1). We therefore tested whether CetZ2 influences cell shape in stationary phase compared to mid-log phase using deletion, overexpression, and GTPase mutations.

Mid-log cultures of these strains were diluted to a theoretical OD_600_ of 0.0005 in 10 mL Hv-YPCab medium with 0.2 or 1 mM ʟ-tryptophan (Trp) to induce expression of the relevant plasmid-encoded genes under the control of the p.*tna* promotor. Samples for mid-log, early stationary, mid-stationary, and late stationary phases were taken after 24, 72, 96, and 120 h of incubation, respectively (see Fig 1a).

At the 24 h mid-log stage, where wild-type cells are largely in a rod-shaped form before reverting to plate shaped cells (1, 21), deletion and overexpression of *cetZ2* had no obvious effect on cell shape, whereas production of the GTPase inactive mutant, CetZ2.E212A, strongly disrupted rod development (Fig. 2a, top panels). These observations were quantified and confirmed by analysis of the circularity of cell outlines (Fig. 2b, left). In late stationary phase (120 h), wild-type cells were a mixture of plate and short rod shapes (Fig. 2a, lower panels). However, deletion of *cetZ2* resulted in more elongation of cells (120 h), whereas overexpression of *cetZ2* or *cetZ2*.E212A increased cell circularity in comparison to the wild-type (Fig. 2a-c). These phenotypes were clear with multiple culture replicates (Fig. 2c) and demonstrate that CetZ2 has a role in maintaining plate shape in stationary phase.

**Figure 2.**
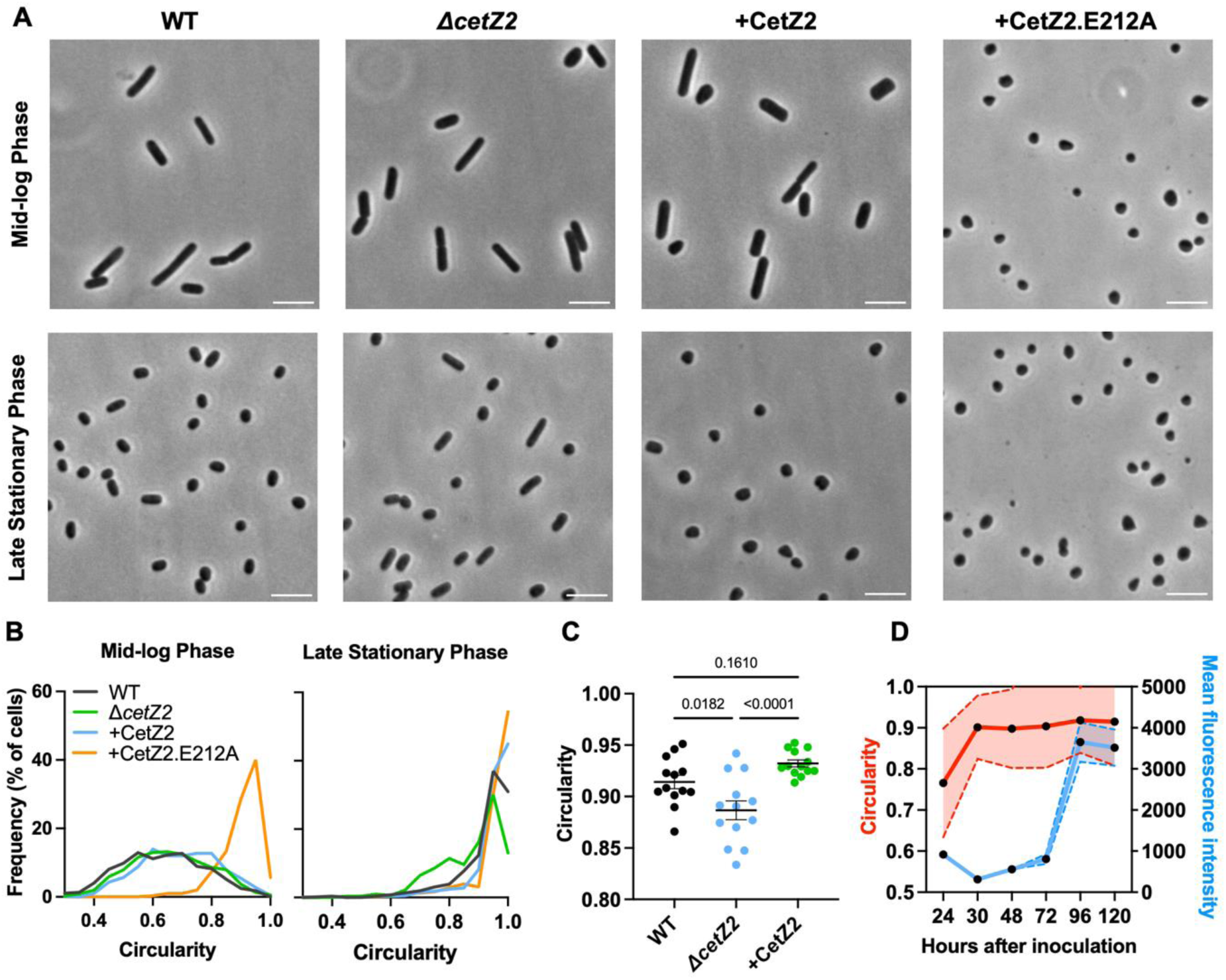
CetZ2 contributes to maintenance of plate-shape in stationary phase. **A)** H26 + pTA962 (WT), Δ*cetZ2* + pTA962, Δ*cetZ2* + pTA962-CetZ2, and WT + pTA962-CetZ2.E212A were grown in Hv-YPCab medium supplemented with 1 mM ʟ-tryptophan and sampled during mid-log phase (24 h after inoculation) and late stationary phase (120 h after inoculation). Phase-contrast microscopy images **(A)** (scale bar=5 μm) and **B)** circularity analysis of individual cells during mid-log and late stationary phase. One representative culture replicate for each strain is shown. **C)** Mean cell circularity (SD) of 13 replicate cultures in late stationary phase (120 h). One-way ANOVA was used as a statistical test. The total number of cells analysed from the 13 culture replicates were: WT (+pTA962), n=14,242, ΔcetZ2 (+pTA962), n=23,671; ΔcetZ2 (+pTA962-cetZ2), n=16,416. **D)** *cetZ2-mTq2CHR* + pTA962 was grown in Hv-YPCab medium from a theoretical starting OD_600_~0.0005 and observed by phase-contrast and fluorescence microscopy after 24, 30, 48, 72, 96, and 120 h of culturing. The circularity and mean fluorescence intensity of individual cells was analysed at each time point. Individual points represent mean of the population at each timepoint, and dotted lines indicate standard deviation. The number of cells analysed at each timepoint is as follows: 24 h, n=253; 30 h, n=554; 48 h, n=1209; 72 h, n=1872; 96 h, n=728; 120 h, n=187.

To investigate the relative timing of CetZ2 upregulation and the transition to plate-shape that occurs in mid-late log growth (19, 25), we developed a chromosomal fluorescent fusion of *cetZ2-mTq2* under the control of the native *cetZ2* promotor (*cetZ2-mTq2CHR*). This strain exhibited cell circularity (Fig. S2a) and CetZ2 upregulation in stationary phase (Fig. S2b) that was equivalent to the wild-type, indicating full functionality of the fusion. The total cellular fluorescence intensity of *cetZ2-mTq2CHR* and cell circularity were then measured for individual cells during the transition to stationary phase (e.g. Fig. S2c). This showed that CetZ2 upregulation occurred between 72 and 96 hours of culturing (Fig. 2d, blue), which was well after the completion the transition of rod-shaped cells to plate-shaped cells at approximately 30 h (Fig. 2d, red). This suggested that CetZ2 upregulation is not linked to mid-log plate shape development, and rather has a role in maintenance of plate-shape in stationary phase.

Previous work showed that *cetZ2* is dispensable for swimming motility (1), but that its overexpression can slightly enhance motility (21). The actively swimming cells at the leading edge of the expanding halo in soft agar motility assays are rod-shaped, whereas those in the interior of the halo (that are slower or not swimming) have transitioned to plate shapes (1). We therefore investigated the relative expression of *cetZ2* in these cells. The *cetZ2-mTq2CHR* strain was fully functional in swimming motility (Fig. S2d, e) and fluorescence measurements indicated that CetZ2-mTq2 was only detectable above background in plate-shaped cells sampled from the inner motility halo (Fig. S2f-i). Thus, the relationship between CetZ2 levels and cell morphology is consistent between batch culture and motility assay conditions, and these observations are consistent with the notion that CetZ2 is only expressed or effective in plate-shaped cells which have already reverted from rod-shape (21).

### CetZ2 subcellular localization patterns change throughout stationary phase

We observed that the *cetZ2-mTq2CHR* strain showed only extremely faint CetZ2-mTq2 subcellular localization relative to the cellular background during stationary phase (Fig. S2c). This was considered likely a combination of the naturally very low concentration of CetZ2 in cells, and the moderate blue auto-fluorescence of *H. volcanii* cells. To enable further investigation of CetZ2 subcellular localization and function, we overexpressed CetZ2-mTq2 or a CetZ2-YPet fusion from a plasmid to achieve a greater signal-to-background ratio. Importantly, both CetZ2-mTq2 and CetZ2-YPet were functional in this context, because they prevented rod formation equivalently to untagged CetZ2 produced from the same vector backbone (Fig. S3a). Likewise, CetZ2.E212A-mTq2 and CetZ2.E212A-YPet had equivalent effects to untagged CetZ2.E212A (Fig. S3b). We therefore expect that any subcellular structures and their behaviours observed under these conditions reflect their function in the prevention of rod formation.

Western blotting confirmed that the plasmid-based system resulted in relatively strong overproduction of the tagged and untagged CetZ2 and CetZ2.E212A compared to endogenous CetZ2 in the wild-type (Fig. S3c), even with no additional inducer, L-tryptophan (Fig. S3d). Interestingly, when CetZ2-mTq2 production was induced with 1 mM of ʟ-tryptophan throughout the growth cycle, the level of CetZ2-mTq2 still increased substantially in stationary phase (Fig. S3e), suggesting the existence of a post-transcriptional or post-translational mechanism for the regulation of CetZ2 levels.

The plasmid-produced CetZ2-mTq2 was then observed throughout the growth cycle (Fig. 3a). During the mid-log phase, CetZ2-mTq2 localized in patches throughout the cell, around the membrane, at mid-cell, and at the cell poles, similar to the previously reported localization of CetZ1 in rod-shaped cells in liquid culture (1, 22) and consistent with the recently reported localization of overproduced CetZ2 in motile rod-shaped cells (21). We interpret these patterns as cytoskeletal structures, though we note that the structure and arrangement of protein polymers within these in vivo assemblies are currently unknown.

**Figure 3.**
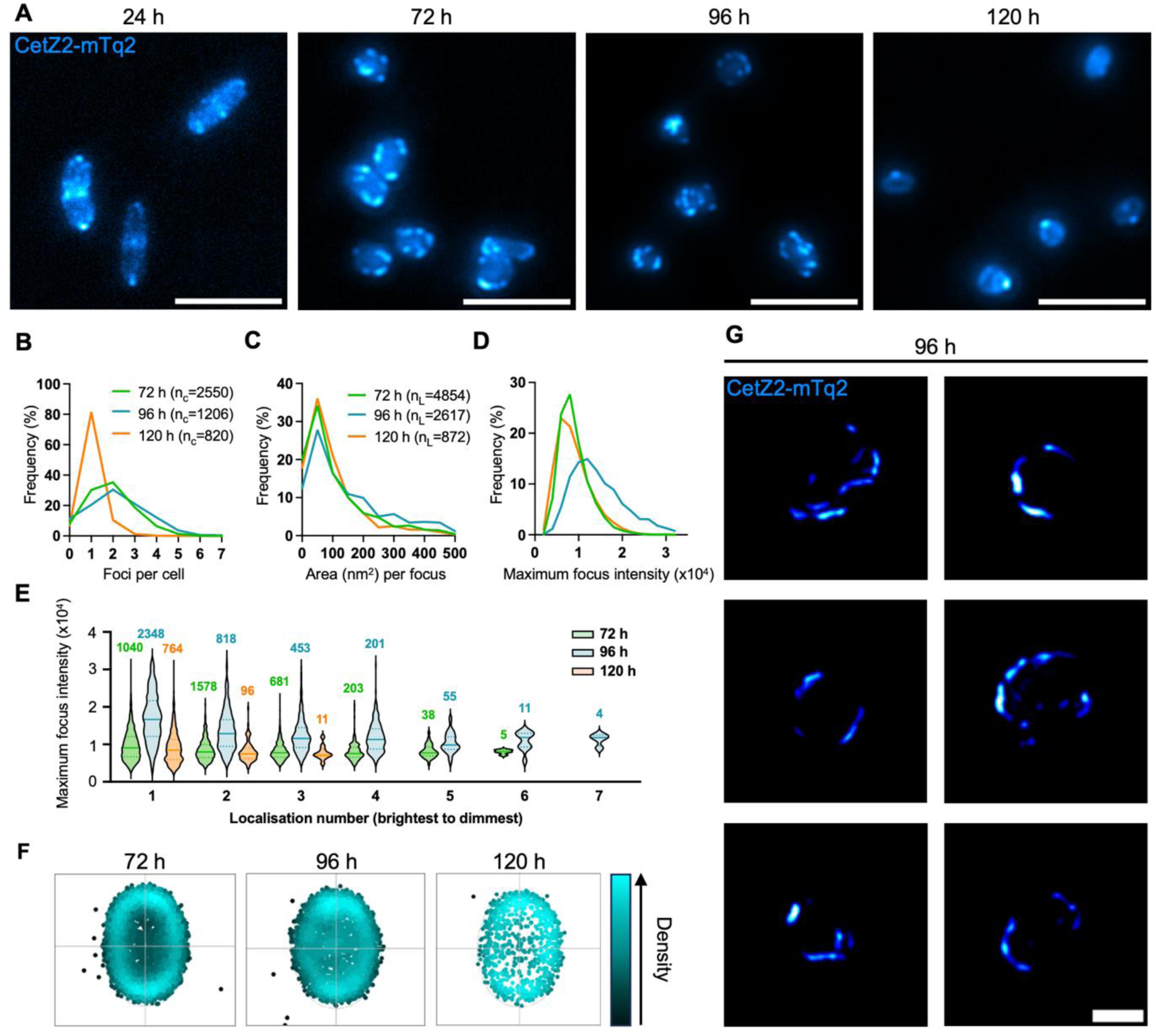
Subcellular localization of plasmid-produced CetZ2-mTq2 in stationary phase. **A)** WT + pHJB6 (for expression of CetZ2-mTq2) was grown in Hv-YPCab medium supplemented with 0.2 mM l-tryptophan and imaged using fluorescence microscopy at 24, 72, 96, and 120 h. Scale bar=5 µm. **B)** 3D-SIM maximum intensity Z-projections of CetZ2-mTq2 at 96 h. Scale bar=2 μm. **C)** CetZ2-mTq2 localization characteristics at 72, 96, and 120 h, including the number of localizations per cell, **D)** Localization area, and **E)** Maximum localization intensity. **F)** Maximum fluorescence intensity by localization number at 72, 96, and 120 h, where localization 1 indicates the brightest detected localization in any given cell, localization 2 indicates the next brightest, etc. The number of localizations constituting each violin plot is detailed above it. **G)** Heatmap representations of CetZ2-mTq2 localizations, coloured by density to indicate frequency of localization. The total number of cells analysed (n_c_) and total number of localizations detected from those cells (n_L_) at each timepoint are indicated in the key in panels **C** and **D**, respectively, and applies to panels **C**-**G**.

In early stationary phase (72 h), CetZ2-mTq2 localized strongly around the cell outline in small patches and filaments. During mid-stationary phase (96 h), CetZ2-mTq2 appeared similar to that in early stationary phase, and an equivalent number of subcellular structures (detected foci) were observed per cell (Fig. 3b). However, localization area (Fig. 3c) and maximum intensity of localizations (Fig. 3d, e) had increased. In late stationary phase (120 h), CetZ2-mTq2 patterns changed substantially, appearing mostly diffuse throughout the cell with one or two cell-edge associated foci (Fig. 3a, b). These foci were decreased in area and maximum intensity in comparison to early and mid-stationary phase, (Fig. 3c-d).

At all timepoints throughout stationary phase, CetZ2-mTq2 was distributed around the cell edge, though showed a weak preference for the polar regions as defined by the longest identified axis (Fig. 3f). To achieve a higher resolution view of these structures, 3D-Structured Illumination Microscopy (3D-SIM) was used to visualize CetZ2-mTq2 during mid-stationary phase (96 h), when localizations were largest and brightest. As may be seen in Fig. 3g and Supplementary Video 1 (SV. 1), this showed clear filaments or patches around the edges of the cell but do not often exist on the top and bottom surfaces. [Note that *H. volcanii* cells have a flattened profile (~0.4 μm thick) and tend to settle on their larger flatter surfaces on agarose pads.]

### CetZ2 cytoskeletal structures undergo dynamic remodelling in mid-stationary phase

Tubulin superfamily proteins, including CetZ1 (1), typically show dynamic remodelling in cells due to their capacity to polymerise and depolymerise in a GTPase-dependent manner, which is central to their cytoskeletal functions. This commonly manifests as polymer treadmilling, where the protein subunits assemble directionally and simultaneously disassemble from the other end. Hence, we used time-lapse imaging of CetZ2-mTq2 to observe any movement of the CetZ2-mTq2 cytoskeletal structures in stationary phase.

In early and late stationary phase cells, imaged over 30 min at 1 min intervals, CetZ2-mTq2 localization appeared quite static, but in mid-stationary phase CetZ2 was relatively dynamic (Fig. 4a, SV. 2). Computational tracking of the filaments and patches (SV. 3) confirmed that CetZ2-mTq2 movement was fastest in mid-stationary phase (Fig. 4b). At this stage, cells displayed three main types of dynamic CetZ2-mTq2 patterns (Fig. 4c, SV. 2). These included (1) a ‘circling’ pattern (42% of cells), in which CetZ2-mTq2 was observed travelling around the perimeter of the cell adjacent to the inner surface of the membrane in a manner consistent with treadmilling, (2) a complex pattern (24% of cells), which showed an apparently more random yet directional motion within the cell, and (3) a ‘confined’ pattern (34% of cells), which were static or relatively slow and envelope associated.

**Figure 4.**
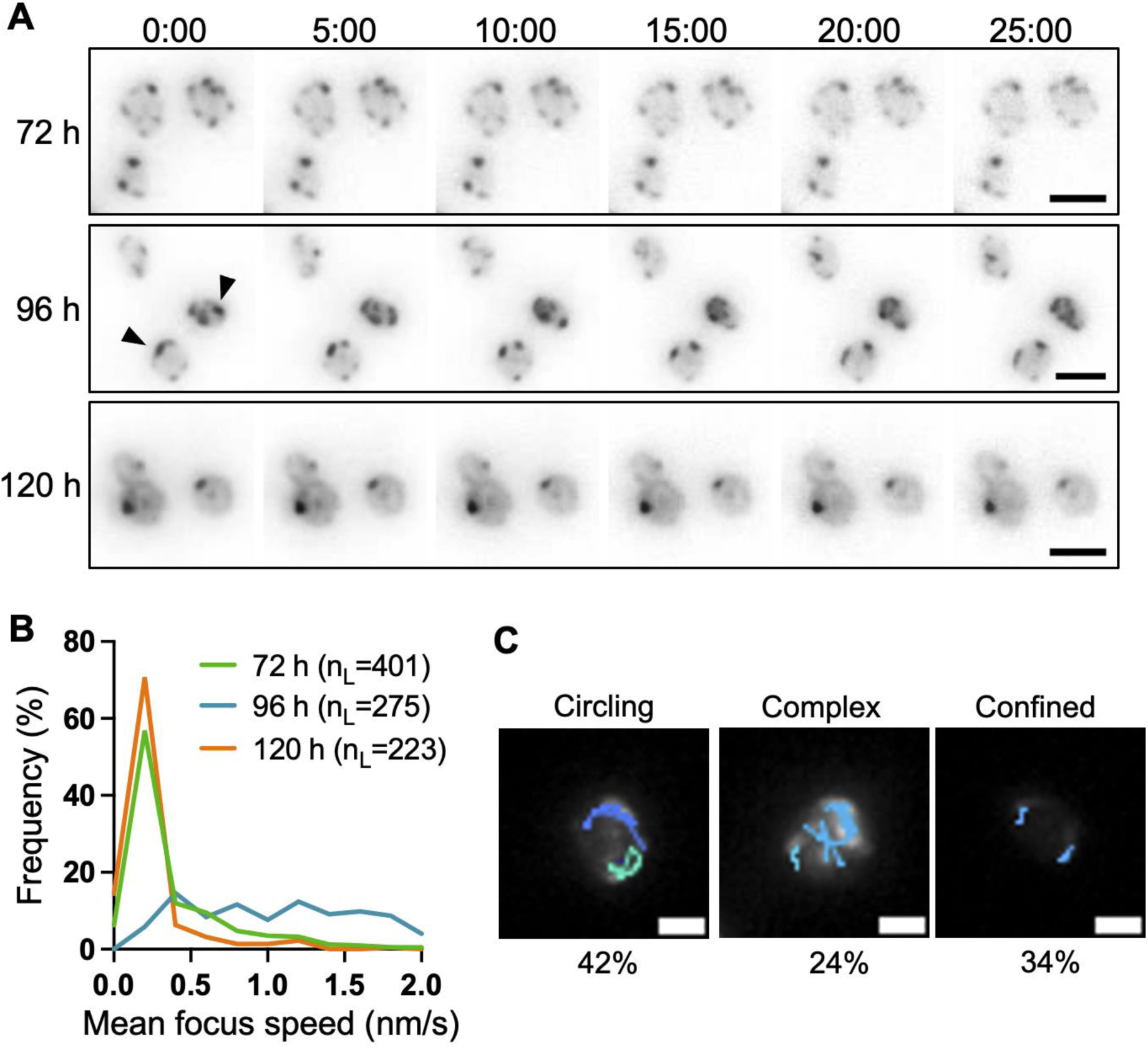
Dynamic behaviours of plasmid-produced CetZ2-mTq2 in mid-stationary phase. WT + pHJB6 cells (for expression of CetZ2-mTq2) were grown in Hv-YPCab medium supplemented with 0.2 mM l-tryptophan and **A)** time-lapse imaging of cells was conducted during early-mid- and late-stationary phase (72, 96, and 120 h, respectively). Images were taken at 1 min intervals for 30 min. Scale bar=2 μm. TrackMate2 was used to track localization speed. Some dynamic structures are showed with arrowheads. **B)** Frequency distribution of localization speed. The total number of detected localizations (n_L_) at each timepoint is indicated in the key. **C)** Overlayed tracks of localization dynamics generated in TrackMate2 were used to define characteristic patterns of CetZ2 dynamics in mid-stationary phase (96 h), determine the proportion of cells displaying each dynamic pattern (n=176). Scale bar=1 μm.

We also investigated the localization and movement of CetZ2.E212A-mTq2 and found that this mutant localized as foci and patches or filaments throughout in cells throughout stationary phase (Fig. S4) but was static at 96 h compared to the wild-type (Fig. S4, SV. 4). Thus, the dynamic structures of CetZ2-mTq2 in stationary phase appear to be dependent on GTPase activity.

### CetZ2 intracellular structures are dependent on the presence of CetZ1

As CetZ2 is only present in species which also have CetZ1 (17), and CetZ2.E212A production inhibits motile rod development (1), it might maintain plate shape in stationary phase by inhibiting CetZ1 function. To begin investigating such potential interactions, we determined whether each protein’s subcellular localization in mid-stationary phase was dependent on the presence of the other. In the absence of CetZ1 (Δ*cetZ1* background), CetZ2-mTq2 did not form patches or large filaments associated with the cell edge as in the wild-type background, but instead mis-localized as discrete and immobile foci that were distributed throughout the cell area (Fig. 5a, g, SV. 5). CetZ2.E212A-mTq2 also mis-localized in the absence of CetZ1, with much fewer foci detected per cell and mostly diffuse fluorescence compared to the strong and stable structures of CetZ2.E212A-mTq2 observed in the wild-type background (Fig. 5b, e, SV. 6).

**Figure 5.**
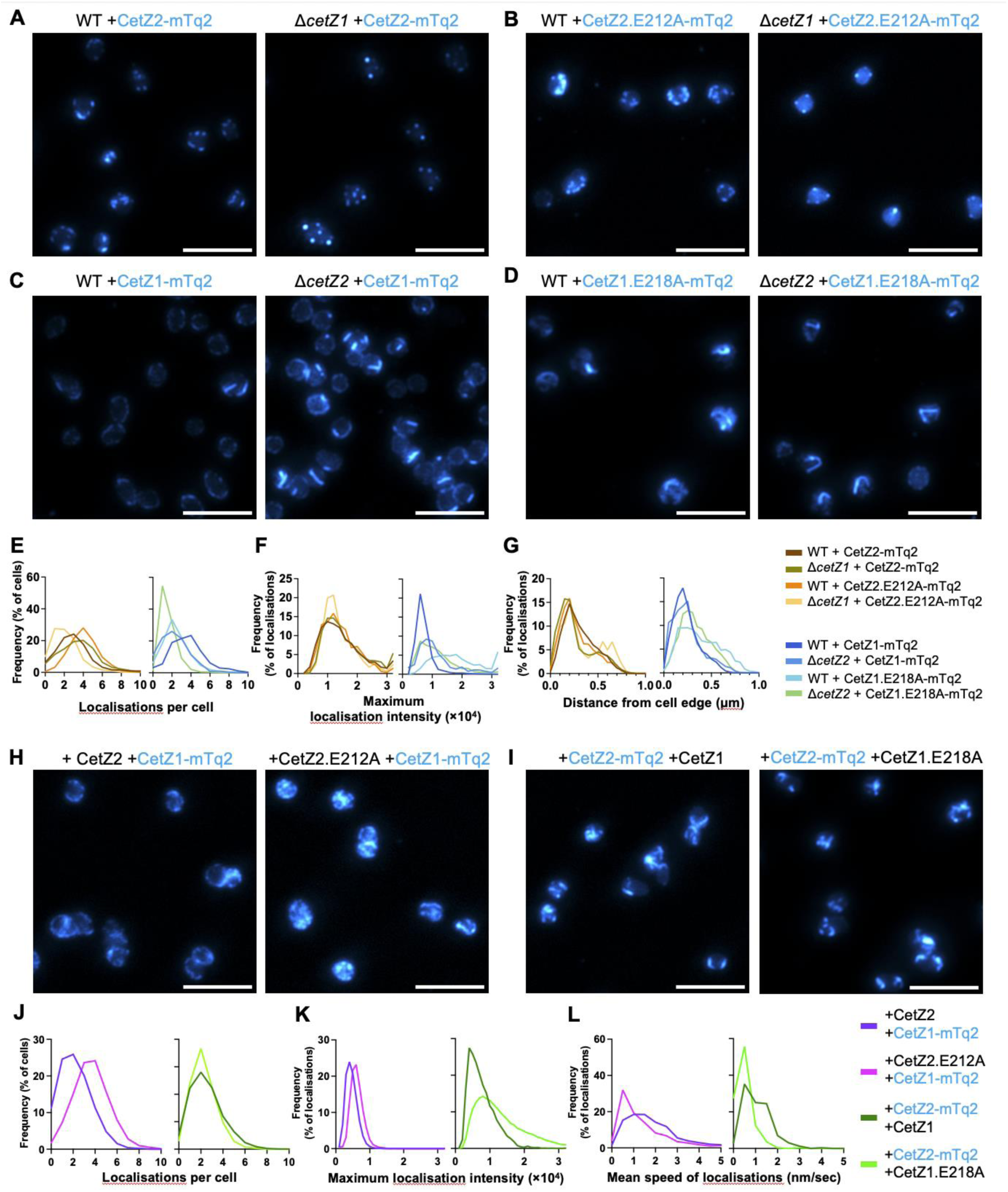
Localization interdependency of CetZ1 and CetZ2 in mid-stationary phase. Cells were grown in Hv-YPCab medium supplemented with 0.2 mM ʟ-tryptophan and imaged after 96 h (mid-stationary phase). **A)** CetZ2-mTq2 (pHJB6) and **B)** CetZ2.E212A (pHJB12) localization in WT and Δ*cetZ1.* **C)** CetZ1-mTq2 (pHVID135) and **D)** CetZ1.E218A-mTq2 (pHJB13) localization in WT and Δ*cetZ2*. For **A**-**D**, scale bar=5 μm. Localization analysis was carried out on this data, measuring **E)** the number of localizations per cell, **F)** maximum intensity of localizations, and **G)** the distance of each localization from the cell edge. The number of cells (n_c_) and number of localizations (n_L_)analysed in **E**-**F** is as follows: WT +CetZ2-mTq2: n_c_=1482, n_L_=4657; Δ*cetZ1* +CetZ2-mTq2: n_c_=904, n_L_=3201; WT +CetZ2.E212A-mTq2: n_c_=347, n_L_=1407; Δ*cetZ1* +CetZ2.E212A-mTq2: n_c_=609, n_L_=1290; WT +CetZ1-mTq2: n_c_=969, n_L_=3558; Δ*cetZ2* +CetZ1-mTq2: n_c_=493, n_L_=1122; WT +CetZ1.E218A-mTq2: n_c_=815, n_L_=1986; Δ*cetZ2* +CetZ1.E218A-mTq2: n_c_=1654, n_L_=2639. **H)** CetZ1-mTq2 was co-expressed with untagged CetZ2 (pHJB43) or untagged CetZ2.E212A (pHJB44) in the WT background. **I)** CetZ2-mTq2 was co-expressed with untagged CetZ1 (pHJB45) or untagged CetZ1.E218A (pHJB46). Localization analysis was conducted on data from **H** and **I**, measuring **J)** the number of localizations per cell and **K)** maximum localization intensity. The number of cells (n_c_) and number of localizations (n_L_) analysed in **J** and **K** is as follows: WT +pHJB43: n_c_=3118, n_L_=16536; WT +pHJB44: n_c_=3960, n_L_=16406; WT +pHJB45: n_c_=2893, n_L_=6256; WT +pHJB46: n_c_=3278, n_L_=56450. **L)** Time-laps imaging was conducted for strains in **H** and **I** (SV. 9 and SV. 10, respectively), and the average localization speed measured. The number of localizations measured was as follows: WT +pHJB43: n=1595; WT +pHJB44: n=524; WT +pHJB45: n=327; WT +pHJB46: n=329. The key in **L** applies to panels **G** and **H**.

During mid-log growth, CetZ2-mTq2 localization patterns were similar to CetZ1-mTq2 (Fig. S5a, b), and the patches were noticeably more distributed throughout the cell area compared to prevalent cell-edge patches seen during stationary phase (Fig. 4a). The mid-log patterns of CetZ2-mTq2 differed in the absence of CetZ1, like we observed during mid-stationary phase (Fig. 5a, S5a). The GTPase mutant CetZ2.E212A-mTq2 partially inhibited rod-shape during mid-log growth when induced using a moderate concentration (0.2 mM) of L-tryptophan, as expected (1), and its subcellular localization also differed in the absence of CetZ1 (Fig. S5c).

### The formation and stability of CetZ1 cytoskeletal structures in stationary phase are dependent on the presence of CetZ2

In mid-stationary phase in the wild-type background, CetZ1-mTq2 localized in patches and small foci, mostly concentrated around the cell edge, but occasionally, larger filaments were also observed (Fig. 5c). Time-lapse imaging showed that while some of the small foci and diffuse regions of CetZ1-mTq2 localization were relatively dynamic, the substantive filaments and patches were stable over the 30 min (SV. 7). The CetZ1-mTq2 localization pattern was similar in the Δ*cetZ2* and wild-type backgrounds (Fig. 5c), but in Δ*cetZ2*, the larger bright filaments were more frequent (Fig. 5c, e, f). Interestingly, in the absence of CetZ2, some of the bright CetZ1 filaments were observed to shrink over the 30 min, which was not apparent in the wild-type background (SV. 7). Together, these results suggest that CetZ2 can lead to inhibition of CetZ1 filament formation yet can stabilize CetZ1 filaments if they form; both effects would be likely to inhibit CetZ1 function.

CetZ1.E218A localized as long, curved, or bent filaments in both the wild-type and Δ*cetZ2* backgrounds (Fig. 5d), but somewhat less frequently and with higher intensity in the absence of CetZ2 (Fig. 5e, f). CetZ1.E218A was not dynamic in either background (SV. 8), demonstrating that CetZ2 is not required for the stabilization mechanism of CetZ1.E218A. During mid-log phase, CetZ1-mTq2 (Fig. S5b) and CetZ1.E218A-mTq2 (Fig. S5d) had indistinguishable localization patterns in wild-type and Δ*cetZ2* backgrounds.

### CetZ1/2 GTPase mutants interfere with the localization dynamics of the other CetZ

Next, we observed the effect of the filament-stabilizing GTPase mutations of CetZ1 or CetZ2 on the localization and dynamics of the other CetZ in mid-stationary phase. This was expected to reveal the extent to which each protein’s GTPase influences dynamic filament turnover of the other protein through potential co-assemblies. Plasmids pHJB43-46 were generated for dual expression of *cetZ2/cetZ1-mTq2*, *cetZ2.*E212A*/cetZ1-mTq2*, *cetZ2-mTq2/cetZ1*, and *cetZ2-mTq2/cetZ1.*E218A, respectively; the *cetZ2* ORFs were located immediately downstream from the p.*tna* promotor, followed by the *cetZ1* ORFs in a bicistronic configuration.

The patterns of CetZ1-mTq2 localization were similar during co-expression with *cetZ2* or *cetZ2.*E212A (Fig. 5h) and showed somewhat larger irregular CetZ1-mTq2 patches and filaments compared to the wild-type and Δ*cetZ2* backgrounds (Fig. 5c). The number of CetZ1-mTq2 localizations per cell was greater during expression with *cetZ2.*E212A compared to *cetZ2* (Fig. 5j), suggesting that CetZ2.E212A may promote or stabilise CetZ1-mTq2 localizations. Time-lapse imaging (SV. 9) and quantification of focus speed (Fig. 5l) revealed that CetZ2.E212A partially stabilized CetZ1-mTq2 localizations.

The patterns of CetZ2-mTq2 localization were similar during co-expression with *cetZ1* or *cetZ1.*E218A (Fig. 5i), and showed somewhat larger irregular CetZ2-mTq2 patches and filaments compared to the wild-type and Δ*cetZ1* backgrounds (Fig. 5a). The number of CetZ2-mTq2 localizations per cell remained similar when co-expressed with *cetZ1* or *cetZ1.E218A*, although *cetZ1.*E218A co-expression resulted in an increase in CetZ2-mTq2 localization brightness (Fig. 5k), and reduced speed (Fig. 5i, SV. 10).

### CetZ1 and CetZ2 localization is partially correlated in mid-stationary phase

The above results suggested that CetZ1 and CetZ2 polymerization-like behaviours are at least partially inter-dependent in stationary phase, consistent with the view that their functions are interlinked, with an antagonistic activity of CetZ2 against the rod development function of CetZ1. This prompted us to determine whether CetZ1 and CetZ2 can colocalize and observe their dynamics in relation to one another during mid-stationary phase. When *cetZ1-*mTq2 and *cetZ2-*YPet were co-expressed from pHJB16 in the wild-type background in mid-stationary phase, both proteins displayed patchy, mostly membrane-associated localizations (Fig. 6a), like their observed localizations when expressed in isolation (Fig. 3a, 5a, c). Co-localization analysis showed that CetZ1-mTq2 and CetZ2-YPet localizations imperfectly overlapped, and the proportion of CetZ2-YPet overlapping with CetZ1-mTq2 (M2=82%) was somewhat higher than the inverse relationship (M1=70%) (Fig. 6d), suggesting there is slightly more ‘free’ CetZ1-mTq2 than CetZ2-YPet in this overexpression experiment. Pearson correlation analysis showed that 36% of cells had a correlation coefficient ≥ 0.7 (Fig. 6e), further demonstrating their significant overlap.

**Figure 6.**
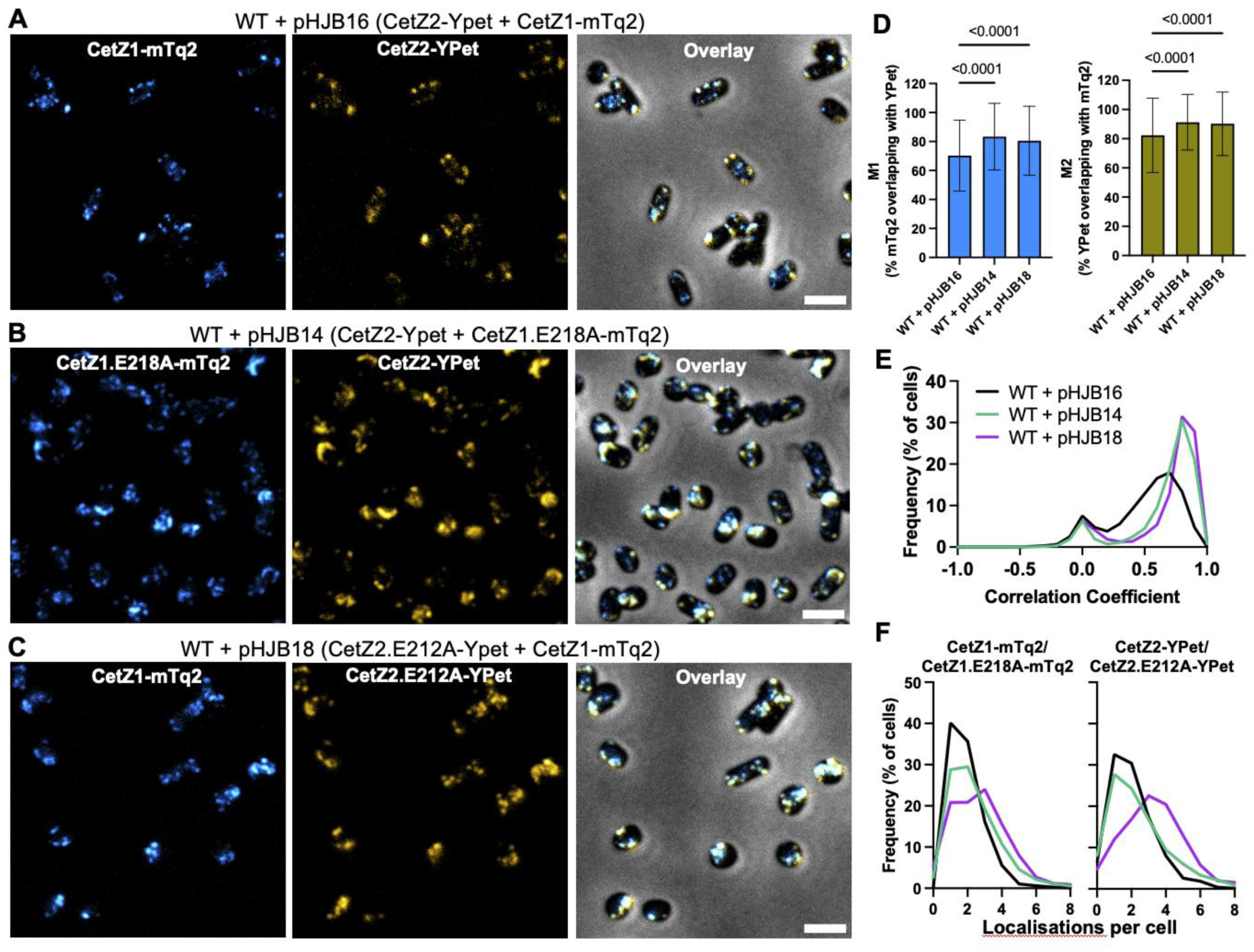
Co-expression and localization of tagged CetZ1 and CetZ2. Wild-type cells expressing CetZ1-mTq2 and CetZ2-YPet (pHJB16), CetZ1.E218A-mTq2 and CetZ2-YPet (pHJB14), or CetZ1-mTq2 and CetZ2.E212A-YPet (pHJB18) were grown in Hv-YPCab medium supplemented with 0.05 mM ʟ-tryptophan to induce expression of fluorescent proteins. **A)** WT + pHJB16, **B)** WT + pHJB14, and **C)** WT + pHJB18 were visualised using phase contrast and fluorescence microscopy during mid-stationary phase. Scale bars=2.5 µm and apply to all panels. Images of each strain were analysed using EzColocalization. **D)** Degree of fluorescent area overlap between mTq2 and YPet as a percentage of total fluorescent area. Mean and standard deviation are shown. One-way ANOVA was used as a statistical test. **E)** Pearson correlation coefficients of CetZ1/CetZ1.E218A-mTq2 and CetZ2/CetZ2.E212A-YPet fluorescence per cell, represented as a distribution. A small peak at 0 was observed for all strains, resultant from a small proportion of cells which only displayed fluorescence in one channel, or no fluorescence. **F)** The number of CetZ1/CetZ1.E218A-mTq2 and CetZ2/CetZ2.E212A-YPet localizations per cell were counted for each strain. The number of analysed cells per strain in **D**-**F** are as follows: WT +pHJB16: n=4643; WT +pHJB14: n=1506; WT +pHJB18: n=1452. Key in panel **E** also refers to panel **F**.

We then co-expressed the two tagged proteins where one was the hyperstable GTPase mutant, to assess whether this would result in greater co-assembly (Fig. 6b, c). The CetZ1.E218A-mTq2 overlap with CetZ2-YPet (M1) increased to 83% while the CetZ2-YPet overlap with CetZ1.E218A-mTq2 (M2) increased 91% (Fig. 6d), and ~70% of cells had a Pearson’s correlation coefficient ≥ 0.7 (Fig. 6e). Indistinguishable results were obtained with the alternate CetZ2.E212A-YPet and CetZ1-mTq2 combination (Fig. 6d, e). Finally, the number of foci per cell moderately increased in the strains carrying a GTPase mutant, compared to the dual wild-type proteins (Fig. 6f). In summary, both combinations with the GTPase mutations showed a substantial increase in the degree of co-localization compared to the two wild-type proteins, demonstrating a strong interdependency between the GTPase functions of the two proteins that affects their direct or indirect co-assembly.

## Discussion

In this study, we describe the function of archaeal tubulin-like protein CetZ2 and its interplay with its paralog CetZ1 that leads to the coordination of shape control and pleomorphism in the growth cycle of the model archaeon *H. volcanii*. In early log phase growth, *H. volcanii* cells largely transition from the plate (or disk-like) cells into rod shaped cells, until later in log phase when they revert into plates and are maintained as such in stationary phase (1, 18). CetZ1 is necessary for plate cells to develop into rods, and it does so independently of CetZ2 (1), which we found here is only produced at a very low level in these conditions. The mid-log reversion from rods to plates is mediated by several other factors including DdfA and the cytoskeletal proteins volactin and halofilins HalA/HalB (26, 27). Here, we saw that CetZ2 does not have a clear role in rod development or reversion to plates, both of which occur well before its strong upregulation as cells enter stationary phase. CetZ1 was much more abundant than CetZ2 throughout the growth cycle, but it is moderately downregulated in stationary phase. The strong increase in the ratio of CetZ2 to CetZ1 as cells enter stationary phase correlated with the observation that CetZ2 is required to help maintain plate shape specifically in stationary phase.

During the peak time of CetZ2 activity in mid-stationary phase, we observed that an overproduced fluorescent-tagged CetZ2 formed clear filaments or patches around the edge of the cells that were noticeably larger and more dynamic over the course of 30 min compared to cells in other phases. The tagged protein showed equivalent function to the overproduced untagged protein in the maintenance of plate cell shape, suggesting that the specific mid-stationary phase patterns reflect functional cytoskeletal structures. Although we cannot rule out the possibility that the necessary overproduction of these proteins, relative to the very low concentration of endogenous CetZ2, may generate an exaggerated pattern, the specificity of these behaviours to a mid-stationary phase argues that they reflect a specific biological role of CetZ2 at this time. Interestingly, the movement of the labelled CetZ2 around the cell was clearly directional, displaying a circling-like behaviour in many cases, which likely reflects protein polymer treadmilling and a cytomotive function of CetZ2 (28); exactly how this could lead to plate shape maintenance is yet to be determined.

The above findings were taken as the basis for investigating potential interplay between CetZ1 and CetZ2 functions, which suggested that CetZ2 contributes to plate shape maintenance in stationary phase by counteracting the CetZ1-based rod-development pathway (schematic in Fig. 7). This manifested as CetZ2-dependent effects on CetZ1 cytoskeletal structures and their dynamic behaviours in stationary phase cells. For example, CetZ1 assemblies were more unstable in the absence of CetZ2 in mid-stationary phase. If CetZ2 directly or indirectly hyper-stabilizes CetZ1 polymers, this would inhibit CetZ1’s rod-development function, which relies on GTPase-dependent disassembly and dynamic turnover of its cytoskeletal structures (1). Secondly, we found that production of the CetZ2 GTPase mutant, which hyper-stabilizes the protein filaments, increased the extent of the observed colocalization of CetZ1 and CetZ2 in stationary phase, further suggesting that the assembly of CetZ1 and CetZ2 cytoskeletal structures are interlinked. Consistent with this, we also observed a significant dependency of CetZ2 cytoskeletal dynamics on the presence of CetZ1 in stationary phase. Together, the results suggest that the two proteins might directly or indirectly interact to counteract the CetZ1-based rod development pathway during stationary phase.

**Figure 7.**
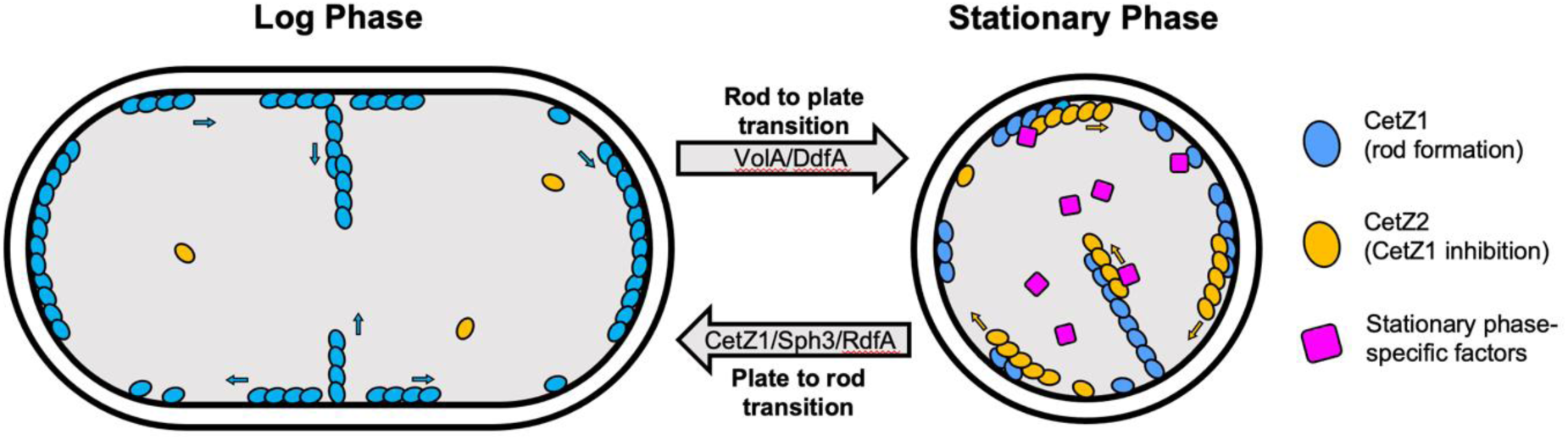
Model for the coordination of shape-change by CetZ proteins in the *H. volcanii* growth cycle. In early and mid-log phase, CetZ1 is abundant and forms dynamic structures adjacent to the inner membrane at cell poles, mid-cell, and the long axis of cells (22), and is necessary for rod-development with other factors such as RdfA and Sph3 (27). CetZ2 is at a very low level and not involved in either of the cell shape transitions in early/mid-log. The rod-to-plate cell transition in the mid-late log phase before cells enter stationary phase is likely controlled by a number of factors such as volactin (VolA) and DdfA (27). In stationary phase (right), CetZ2 is upregulated, and CetZ1 is downregulated, which might enable CetZ2 to counteract CetZ1 function, through direct or indirect interactions, thereby maintaining plate-shape by preventing rod-development in stationary phase.

Interestingly, the effects of CetZ2 on preventing rod formation were highly specific to stationary-phase. Overproduction of CetZ2 was only capable of inhibiting rod shape in stationary phase and not mid-log phase. Similarly, CetZ1 cytoskeletal structures were only affected by CetZ2 overproduction in stationary phase. We therefore believe it is likely that there are other factors specifically present in stationary phase that are necessary for CetZ2-based maintenance of plate cell shape (Fig. 7, right). The identity of these potential factors is currently unknown, but the involvement of other proteins would be unsurprising, given tubulin and FtsZ both recruit and work in conjunction with a myriad of other proteins (10, 29–32).

The other four *H. volcanii* CetZs (CetZ3-6) might represent candidates. However, previous proteomic studies did not detect upregulation of these in stationary phase (24). Although this does not rule out potential roles for them with CetZ2 in plate cell maintenance, it suggests they are not likely to be the stationary-phase specific factors implicated by our findings. CetZ5 has been hypothesised to be involved in the rod-to-plate transition based on proteomics that compared rods from early log and plates from late log (27), but this potential role has not yet been experimentally tested, and would nevertheless occur well in advance of the mid-stationary phase functions characterized here. Further genetic analyses or protein association and identification studies are hence warranted to begin characterizing the detailed mechanisms underpinning the stationary-phase cell shape control that we have discovered here.

In Haloarchaea, CetZ1 and CetZ2 homologs group into distinct protein families amongst multiple additional CetZ homologs in some species that are not so clearly grouped (17). Interestingly, CetZ2 was only identified in haloarchaea that also had CetZ1 (17). This raised the hypothesis that there is a functional dependency of CetZ2 on CetZ1, as identified here. Since most haloarchaea with both CetZ1 and CetZ2 are pleomorphic and rod forming, while those that had CetZ1 but not CetZ2 were not pleomorphic (17), the mechanism of coordination of shapeshifting though CetZ1 and CetZ2 is likely to be conserved across other Haloarchaea.

Our findings demonstrate that CetZ paralogues in haloarchaea can possess distinct and opposing functions, consistent with the phylogenetic separation and conservation of their two CetZ paralogous groups (1, 16, 17). The diversification of CetZ1 and CetZ2 function and their coordination is reminiscent of the specialised and coordinated functions of FtsZ1 and FtsZ2 in *H. volcanii* cell division (33) and of the multiplication, diversification, and specialisation of tubulins in eukaryotes. Functional antagonism is rare amongst cytoskeletal proteins, but notably is a function of the vertebrate β5-tubulin, which destabilizes microtubules when upregulated (34). We speculate that the opposing functions of archaeal CetZ1 and CetZ2 might resemble an early step towards the evolution of obligate hetero-polymerisation, such as seen between the active and inactive α- and β-tubulins in microtubules of modern-day eukaryotes.

## Materials and Methods

### Materials and reagents

Unless otherwise specified, all chemicals were obtained from Sigma, reagents for cloning procedures were obtained from New England Biolabs and used according to the manufacturer’s instructions, and custom oligonucleotides were obtained from Integrated DNA Technologies. The Meridian Bioscience ISOLATE II PCR and Gel Kit and ISOLATE II Plasmid Mini Kit were used for purification of DNA from PCR reactions and agarose gels, and for plasmid extractions, respectively.

### *H. volcanii* strains and growth

A list of *H. volcanii* strains used in this study is available in Table S1. Cultures were prepared where indicated in standard culture media, either Hv-Cab (1), Hv-YPCab (18), or Hv-Min with trace elements (35) and incubated at 42 °C with rotary shaking (200 rpm). If necessary, medium was supplemented with uracil (50 µg/mL) to fulfil auxotrophic requirements of Δ*pyrE2* strains, or with ʟ-tryptophan to induce expression of genes under the control of the *p.tna* promotor. For experiments that compared mid-log and stationary phase, cells were cultured in 10 mL Hv-YPCab medium by dilution of a steady mid-log starter culture to a theoretical starting OD_600_ of 0.0005. Mid-log phase samples of cells were taken after a further 24 h of culturing, and then for early, mid, and late stationary phase, samples were taken at 72, 96, and 120 h, respectively.

### Genetic modification

The wild-type H26 (ID621) genetic background of *H. volcanii*, which is auxotrophic for uracil (Δ*pyrE2*), was used throughout this study. To generate the *cetZ1* and *cetZ2* knockout strains, pTA131-based plasmids for deletion of *cetZ1*(18) and *cetZ2*(1), containing the upstream and downstream flanks of *cetZ1* and *cetZ2* and the *pyrE2* gene, were used to transform H26 with selection on Hv-Cab agar. Transformation was done using the previously described method (36) and two-step (‘pop-in-pop-out’) homologous recombination (37) was used for selection of transformants that showed complete deletion of the target gene.

To generate the *cetZ2-mTq2CHR* chromosome tag strain, pTA131-*cetZ2mTq2* was obtained synthesised (Genscript), which contained a 350 bp fragment encoding the C-terminal portion of *cetZ2*, fused in frame to an ORF containing the EG linker (semi-rigid) and mTurquoise2 (20), followed by the 350 bp flank immediately downstream of the *cetZ2* stop codon. This plasmid was used to transform H26 by the two-step method (37) to produce a scarless insertion of EG-mTurquiose2 before the native *cetZ2* stop codon.

### Construction of plasmids for gene expression in H. volcanii

A list of plasmids used in this study is available in Table S2 and relevant oligonucleotides are listed in Table S3. To generate plasmids for single and dual expression of CetZ fusion proteins, we used a recently described plasmid-based expression system (20). Briefly, the ORFs for *cetZ2* or *cetZ2.*E212A excluding their stop codons were PCR amplified using primers to incorporate NdeI and BamHI sites at the 5’ and 3’ ends, respectively, and the resulting PCR product and the vector backbone were ligated using T4 DNA ligase. The vector backbone was pHVID21 for C- terminal mTurquoise2 fusions with the EG linker, or pHVID20 for C-terminal YPet fusions with the EG linker.

To generate plasmids for dual expression of two tagged proteins, pHJB5 and pHJB11 (containing *cetZ2*-YPet and *cetZ2*.E212A-YPet, respectively) were used as vector backbones for insertion of *cetZ1*-mTq2 and *cetZ1*.E218A-mTq2 ORFs positioned immediately downstream of the *cetZ2* ORFs as described (20). The ORFs for *cetZ2-*YPet, *cetZ2.*E212A-YPet, *cetZ1*-mTq2, and *cetZ1.*E218A*-*mTq2 were sourced from pHJB5, pHJB11, pHVID135, and pHJB13, respectively.

Plasmids for dual expression of one untagged CetZ and one tagged CetZ were prepared using plasmids for expression of *cetZ2*-mTq2, *cetZ2*, or *cetZ2*.E212A (pHJB6, pTA962-*cetZ2*, and pTA962-*cetZ2*.E212A, respectively) as backbones. pHJB6 was subjected to restriction digest by NheI and NotI, while pTA962-*cetZ2* and pTA962-*cetZ2*.E212A were digested using BamHI and NotI. ORFs for *cetZ1* and *cetZ1.*E218A were amplified by PCR incorporating an NheI site at the 5’ end, and the ORF for *cetZ1*-mTq2 was amplified to incorporate a BglII site at the 5’ end. After restriction digest with the relevant enzymes, the various *cetZ1* ORFs were ligated as described above, into the backbones, generating plasmids pHJB43-46 for dual expression of *cetZ2/cetZ1-*mTq2, *cetZ2.*E212A/*cetZ1*-mTq2, *cetZ2-*mTq2/*cetZ1*, and *cetZ2-*mTq2/*cetZ1.*E218A, respectively. All plasmid regions originating from PCR products were sequenced verified (Australian Genome Research Facility). Plasmids were passaged through *E. coli* strain C2925 for demethylation before transformation of *H. volcanii*.

### Soft-agar motility assay

*H. volcanii* was maintained in exponential growth in liquid culture (Hv-Cab medium) for at least two days. Following this, cultures were normalised to OD_600_~0.1, and 2 µL of cells were spotted onto the surface of Hv-Cab agar (0.2% w/v). Soft-agar plates were incubated statically at 42 °C for 4 days before extracting cells from the halo edge or inner halo (approximately 5 mm from the inoculation site) as described (21) for further analysis by microscopy.

### Microscopy

For all microscopy, 2 µL of cells were spotted onto a 1.5% (w/v) agarose pad containing 18% buffered salt water (37), and a glass coverslip (#1.5) was placed on top before imaging. Phase contrast and fluorescent images were acquired using a V3 DeltaVision Elite inverted fluorescence microscope (GE Healthcare), with CFP (excitation filter = 400-453 nm; emission filter = 463-487 nm) and YFP (excitation filter = 497-527 nm; emission filter = 537-549 nm) filter sets. Images were acquired using a 100X 1.4 NA Plan Apo objective. Time-lapse images were recorded over 30 min with 1 min intervals, using an incubated microscope stage at 37 °C. Exposure and acquisition settings were maintained between strains and replicates within each experiment.

Three-dimensional structured-illumination microscopy (3D-SIM) was carried out as previously described (22, 23), using a DeltaVision OMX SR (GE Healthcare) microscope with a 60X Plan 1.42 NA objective, a PCO sCMOS camera, 125 nm Z-step size, with a DAPI excitation filter (405 nm) and AF488 emission filter (504-552 nm). Raw images were reconstructed using SoftWorX (Applied Precision, GE), and then maximum intensity projections were generated in FIJI (38), and 3D renders were generated using Imaris (V 7.6.4 BitPlane Scientific).

### Image analysis

All shape analysis of cells was conducted using phase-contrast images analysed using MicrobeJ v5.13m (39) in FIJI (38). To detect cell outlines using phase-contrast images default settings were applied whilst restricting area between 0.8-max µm^2^, and sinuosity between 0-2. For all analysis of fluorescence, raw images and time-lapse data were first processed by applying the *Background Subtraction* function (rolling ball size = 8 pixels). Time-lapse data were further processed using *Bleach Correction* (algorithm = *Histogram Matching*), and *StackReg* (transformation = *Translation*).

For static localization analysis, the *foci* detection method with default settings was used with tolerance set to 1000, Z-score was set to 25, and area was restricted between 0-0.5 µm^2^. Mean whole-cell fluorescence intensity (Bacteria.Intensity.ch2.mean), the number of localizations per cell (Bacteria.Maxima), localization area (Maxima.Shape.area), and maximum localization intensity (Maxima.Intensity.max) were extracted from MicrobeJ results tables, and localization heat maps were generated using the *XYCellDensity* function. For tracking of localizations, TrackMate2 v7.7.2 (40, 41) was used. The LoG detector was used to detect foci, using an estimated foci diameter of 0.5 µm and a quality threshold of 100. For tracking, the LAP tracker was used, allowing for frame-to-frame linking, track segment gap closing, track segment splitting, and track segment merging, with a maximum distance of 0.5 µm and a maximum frame gap of 2.

EzColocalization (42) was used for colocalization analysis. Manual adjustment of thresholds using the default algorithm was required for accurate detection of fluorescent channels, ensuring that weak or background fluorescence was not included in the thresholding. Auto thresholding with the default algorithm and watershed segmentation was used for cell identification using the phase-contrast channel. Fluorescent channels were aligned, and the background was subtracted using a rolling-ball size of 8. All remaining settings and parameters were used as default.

### SDS-PAGE and Western blotting

Samples were prepared for SDS-PAGE by resuspending whole cell pellets in lysis buffer (20 mM Tris-HCl pH 7.5, 1X DNAseI (ThermoFisher), 1X EDTA-free protease inhibitor (ThermoFisher)) and incubating for 30 min at room temperature. Following that, 4X SDS-PAGE sample buffer (250 mM Tris-HCl pH 6.8, 40% (v/v) glycerol, 4% (w/v) SDS, 0.04% (w/v) Bromophenol blue, 20% (v/v) 2-mercaptoethanol) was added to a final concentration of 1X, using a volume that would give a theoretical OD_600_ of 5 based on the original culture. The samples heated at 95 °C for 5 min then vortexed for 1 min.

CetZ1-6xHis and CetZ2-6xHis proteins (1)were purified and quantified by A_280 nm_ in a previous study (1) and diluted in 1X SDS-PAGE sample buffer to 200 ng/μL and 50 ng/μL, respectively. Following this, 1:2 serial dilution in 1X SDS-PAGE sample buffer were carried out for each protein. Samples were loaded onto a 4-20% Mini-PROTEAN® TGX™ pre-cast polyacrylamide gel (Bio-Rad), using the Broad Range Blue Prestained Protein Standard (Νew England Biolabs). Electrophoresis was carried out at 100 V for 85 min with a Bio-Rad Mini-Protean3 electrophoresis system.

For western blotting, proteins separated by SDS-PAGE were transferred to a 0.2 µm nitrocellulose membrane using the Trans-Blot Turbo mini transfer system (Bio-Rad). The membrane was stained with Ponceau S (0.1% (w/v) Ponceau S in 5% (v/v) acetic acid) for 5 min and the excess stain removed by rinsing with Milli-Q water to obtain a total protein stain. Following this, the nitrocellulose membrane was blocked in 5% (w/v) skim milk powder in TBST (50 mM Tris-HCl, pH 7.4, 150 mM NaCl, 0.05% (v/v) Tween-20), for 2 h at room temperature. Primary antibodies(1) against CetZ1 or CetZ2 were diluted 1/500 in TBST for incubation with the membrane overnight at 4 °C. The membrane was washed 3 times in TBST and incubated with the secondary antibody (donkey anti-rabbit IgG HRP conjugate, AbCam 16284) diluted 1/5000 in TBST for 2 h at room temperature. The membrane was washed 5 times in TBST and incubated with the SuperSignal™ West Pico PLUS chemiluminescent substrate (ThermoFisher) for 5 min before imaging of the membrane using an Amersham™ Imager 600 (GE Healthcare).

To calculate the number of molecules of CetZ1/2 per cell, six replicate cultures were grown in Hv-YPCab medium and sampled after 24 and 120 h. Band intensity from western blotting analysis (Fig. 1d, e) was determined by measuring the raw integrated density in ImageJ(38) and subtracting the raw integrated density of the background. The same measurements were conducted for pure protein standards of known concentrations. Band intensities of the standards were used to determine the amount of protein per OD unit of cells in mid-log (24 h) and stationary phase samples. The number of cells per OD unit in mid-log and stationary phase was calculated using serial dilutions and counting the number colony forming units from the same six culture replicates used for western blotting samples. Finally, this was used to determine the number of CetZ1 and CetZ2 molecules per cell.

## Acknowledgements

The authors would like to acknowledge L. Cole and A. Bottomley of the Microbial Imaging Facility at UTS for their training, maintenance, and management of the microscopes. The work was supported by the Australian Research Council (DP160101076).

## Author Contributions

HJB and IGD designed the project, experiments, and experimental methodologies, and wrote the manuscript. HJB performed the experiments and data collection, investigation, and formal analysis.

