## Supplementary Information for "Archaeal tubulin-like proteins CetZ1 and CetZ2 have opposing effects on cell morphology during the growth cycle of *Haloferax volcanii*"

### CONTENTS:

#### Supplementary Figures:

1. CetZ1 and CetZ2 expression levels are not dependent on one another.
2. Function and expression of *cetZ2-mTq2* from the chromosome during stationary phase transition and motility assays.
3. Analysis of plasmid-based expression of *cetZ2* or CetZ2 fusion proteins under various experimental conditions.
4. A mutation in the CetZ2 GTPase active site alters subcellular localization and blocks dynamic movement in stationary phase.
5. Localisation of CetZ1, CetZ2, CetZ1.E218A, and CetZ2.E212A during mid-log growth.

#### Supplementary Tables:

1. Strains used in this study
2. Plasmids used in this study
3. Oligonucleotides used in this study

#### Legends for Supplementary Videos

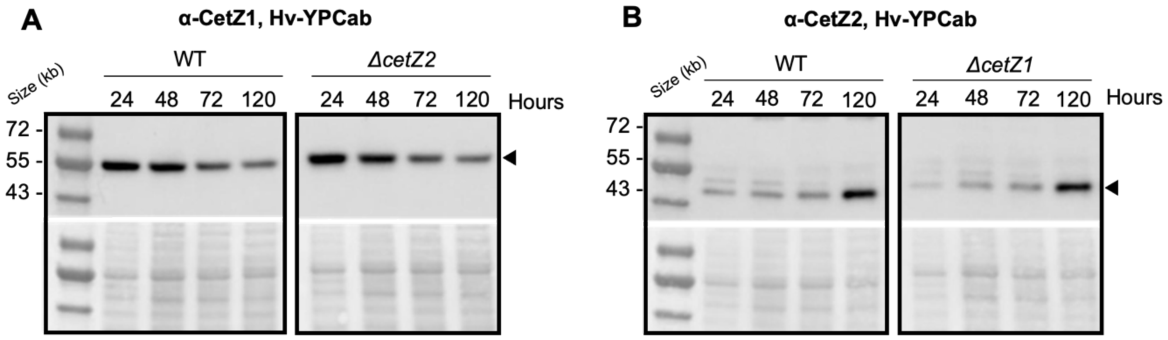


**Supplementary Figure S1.** ***CetZ1 and CetZ2 expression levels are not dependent on one another.*** **A)** Cultures of wild-type and the *cetZ2* deletion strain containing pTA962 were grown in Hv-YPCab medium and whole cell lysate samples were taken after 24, 48, 72, and 120 hours of culturing for western blotting analysis using an antibody against CetZ1. Solid arrowhead indicates band for CetZ1, and bottom panel shows Ponceau S staining used to confirm equal loading of samples. **B)** As in A, but for analysis of CetZ2 expression in the wild-type and *cetZ1* deletion background. Solid arrowhead indicates band for CetZ2.

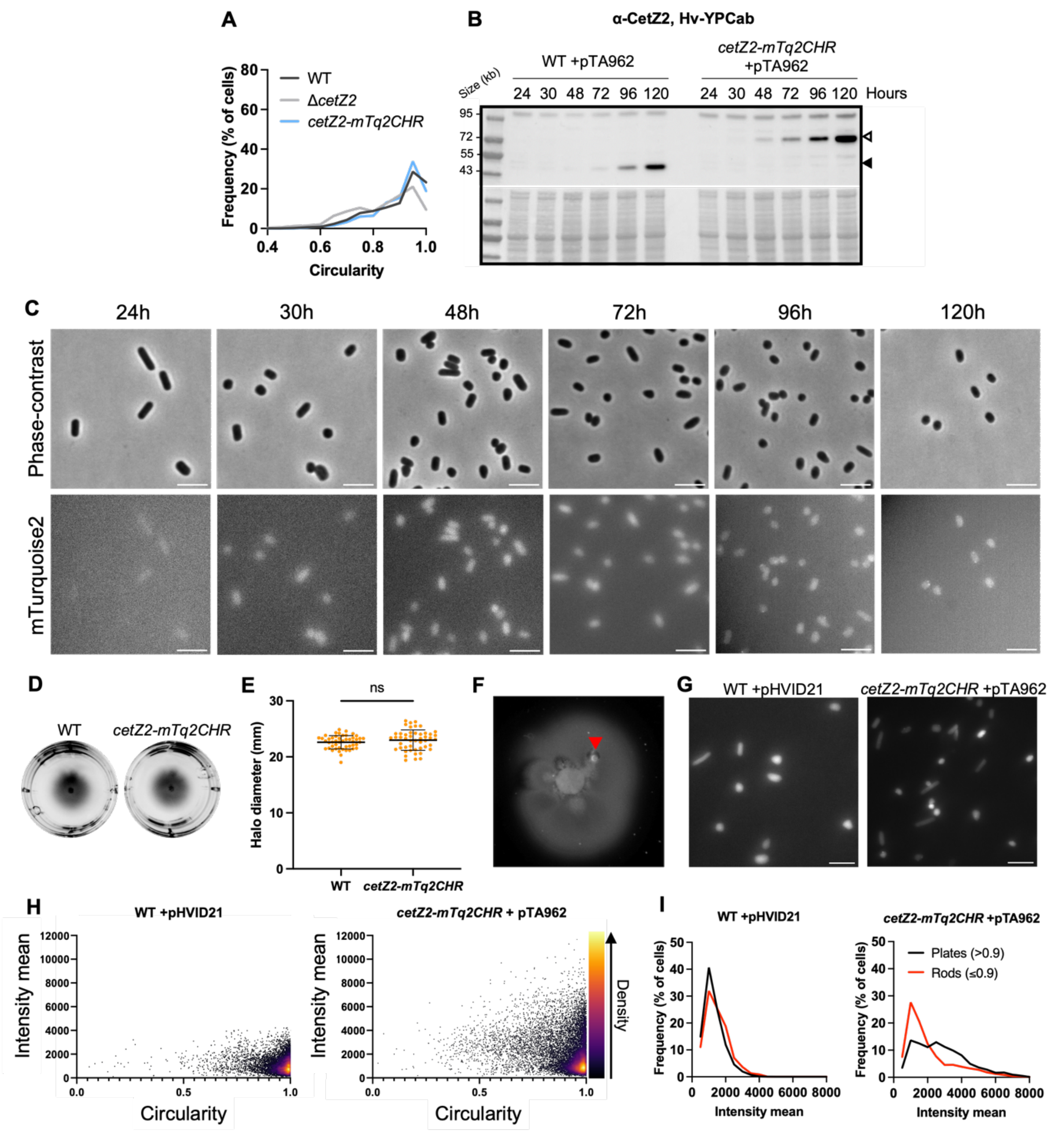


**Supplementary Figure S2.** ***Function and expression of cetZ2-mTq2 from the chromosome during stationary phase transition and motility assays.*** **A)** Cell circularity measurements were obtained from phase-contrast imaging of *cetZ2-mTq2CHR* +pTA962 grown in Hv-YPCab medium supplemented with 0.2 mM L-tryptophan and sampled in late stationary phase (120 h). **B)** WT +pTA962 and *cetZ2-mTq2CHR* +pTA962 were grown in Hv-YPCab and whole cell lysate samples were taken at 24, 30, 48, 72, 96, and 120 h of culturing for western blotting analysis using an antibody against CetZ2. Solid arrowhead indicates band for CetZ2, empty arrowhead indicates band for the CetZ2-mTq2 fusion protein. **C)** Representative phase-contrast and fluorescence microscopy images of *cetZ2-mTq2CHR* + pTA962 grown in Hv-YPCab liquid medium throughout the growth cycle, used to generate graph in Fig. 2D. Scale bar=5 µm. **D)** WT +pTA962 and *cetZ2-mTq2CHR* +pTA962 were inoculated on Hv-Cab soft-agar (0.2%), and incubated for 3 days at 42℃. **E)** The diameter of motility halos was measured for eight culture replicates with 6 technical replicates each. Unpaired t-test was used for statistical analysis, ns=not significant. **F)** WT +pHVID21 (for expression of free mTq2) and *cetZ2-mTq2CHR* +pTA962 were inoculated on soft-agar containing 1 mM L-Tryptophan as in **D**, and were extracted from the inner halo region, approximately 5 mm from the inoculation site after four days. Red arrowhead indicates sample location at inner halo. **F)** Extracted cells were observed by fluorescence microscopy. Scale bar=5 μm. **H)** The average whole-cell fluorescence intensity and circularity (measured using phase-contrast images, not shown) of individual cells from **G** are displayed in XY scatter plots. Individual data points are coloured by density. **I)** Cells with a circularity >0.9 were categorised as plates, and those with circularity ≤0.9 were categorised as rods. The mean fluorescence intensity in plates and rods is represented as a distribution. Data in **H** and **I** was pooled from four biological replicates each with at least two technical replicates. WT + pHVID21: n=7468; *cetZ2mTq2CHR* + pTA962: n=13400.

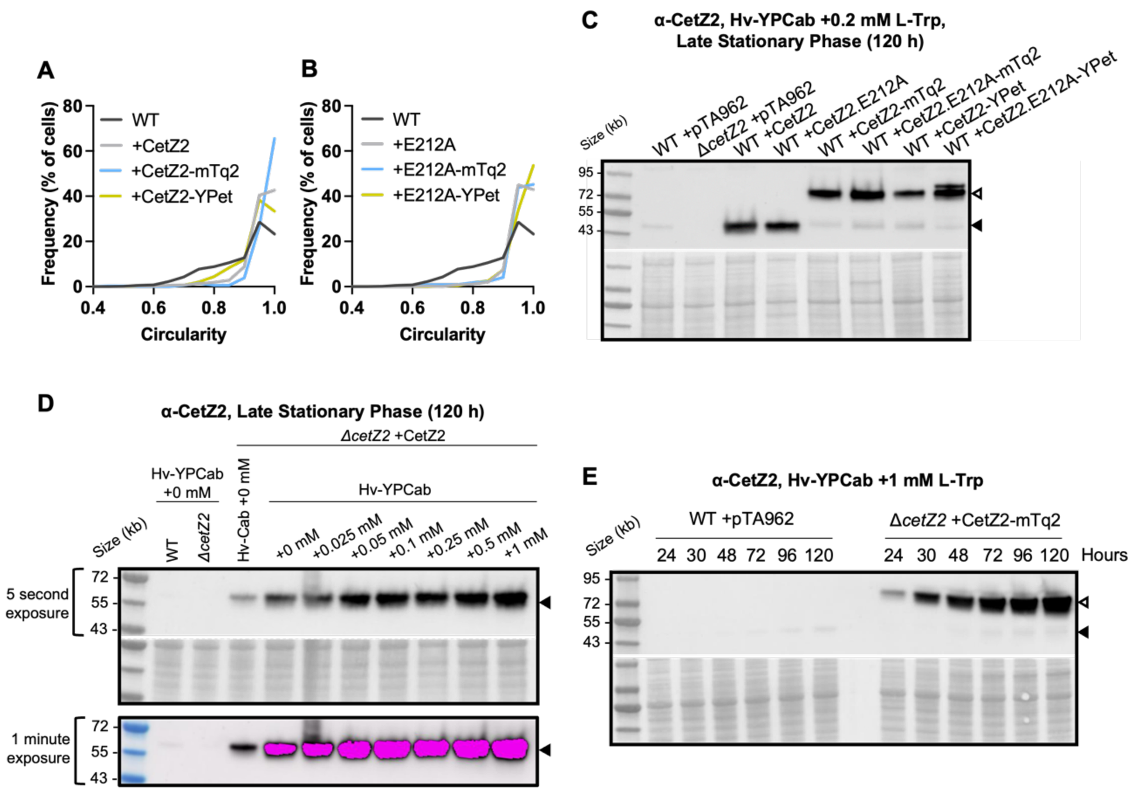


**Supplementary Figure S3.** ***Analysis of plasmid-based expression of cetZ2 or cetZ2 fusion proteins under various experimental conditions.*** **A)** CetZ2 or CetZ2 fusion proteins were produced in Δ*cetZ2* using 0.2 mM L-tryptophan and were sampled during late stationary phase and for phase-contrast microscopy (not shown). The cell circularity for each strain was measured to determine CetZ2 fusion protein functionality in comparison to expression of untagged CetZ2 from the same vector backbone. **B)** As in **A,** but for CetZ2.E212A and CetZ2.E212A fusion proteins. **C)** The CetZ2 and CetZ2.E212A fusion proteins in **A** and **B** were produced in the wild-type (H26) background by induction with 0.2 mM L-tryptophan, and whole cell lysates were taken during late stationary phase (120 h). These were analysed by western blotting to examine proteolytic cleavage. Since induction of CetZ2 or CetZ2 fusion protein production with 0.2 mM L-tryptophan resulted in strong overproduction in **C** and caused an overexpression-like phenotype (increased circularity) in **A**, we sought to compare varying levels of L-tryptophan induction to the native level of CetZ2. **D)** The wild-type (H26 +pTA962) or Δ*cetZ2* (+pTA962) were grown in Hv-Cab or Hv-YPCab medium with the indicated concentrations of L-tryptophan and whole cell lysates were sampled during late stationary phase for western blotting. Longer exposure showed a band for wild-type in Hv-YPCab +0 mM L-tryptophan (lane 2) which was not visible in the shorter exposure, however longer exposure resulted in saturation of chemiluminescent signal from other lanes (indicated in pink). **F)** Wild-type (H26 +pTA962) and Δ*cetZ2* producing CetZ2-mTq2 from pHJB6 were grown in Hv-YPCab supplemented with 1 mM L-Tryptophan and sampled for western blotting analysis at the indicated timepoints throughout the growth cycle. In **C**-**F**, all blots have used an antibody against CetZ2. Solid arrowheads indicate size for CetZ2, empty arrowheads indicate size for CetZ2 fusion proteins. Bottom panels show Ponceau S total protein staining.

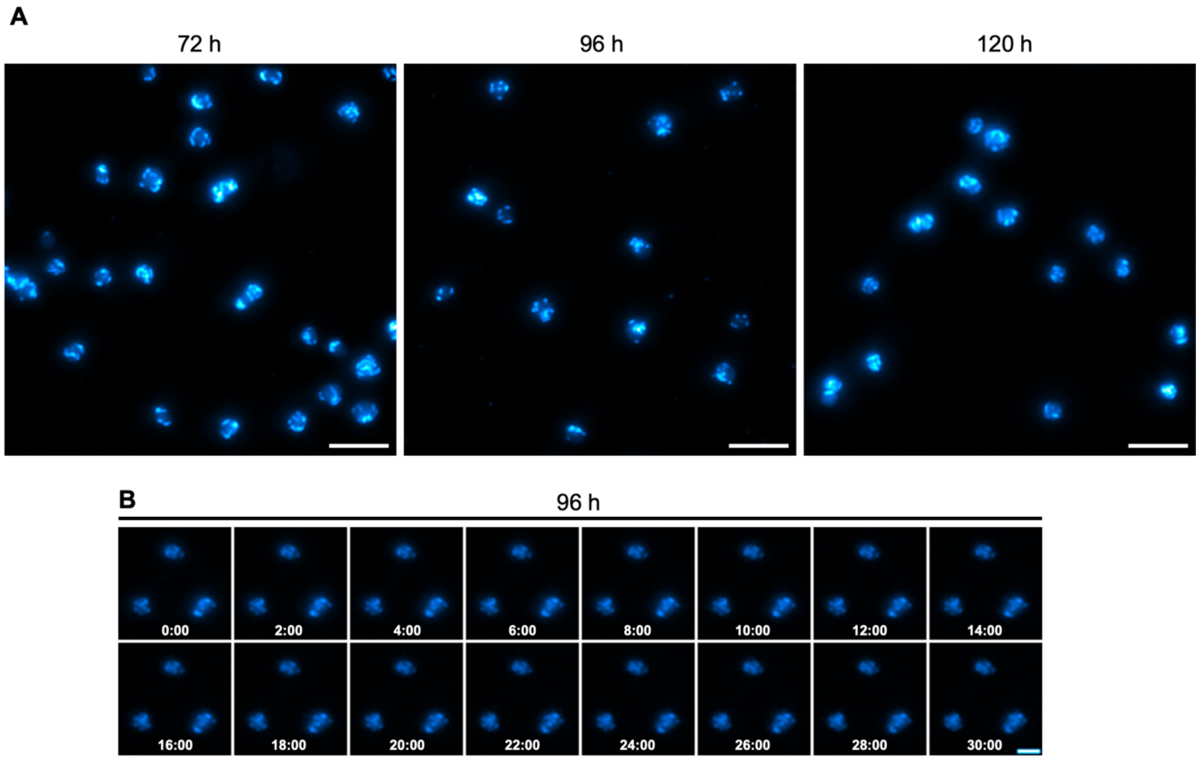


**Supplementary Figure S4.** ***A mutation in the CetZ2 GTPase active site alters subcellular localization and blocks dynamic movement in stationary phase.*** **A)** WT + pHJB12 (CetZ2.E212A-mTq2) was grown in Hv-YPCab medium supplemented with 0.2 mM ʟ-tryptophan and imaged at 72, 96, and 120 h. Scale bar=5 µm. **B)** Time-lapse imaging of cells after 96 h. Images were taken at 1 min intervals for 30 min (SV. 4). Scale bar=2 µm.

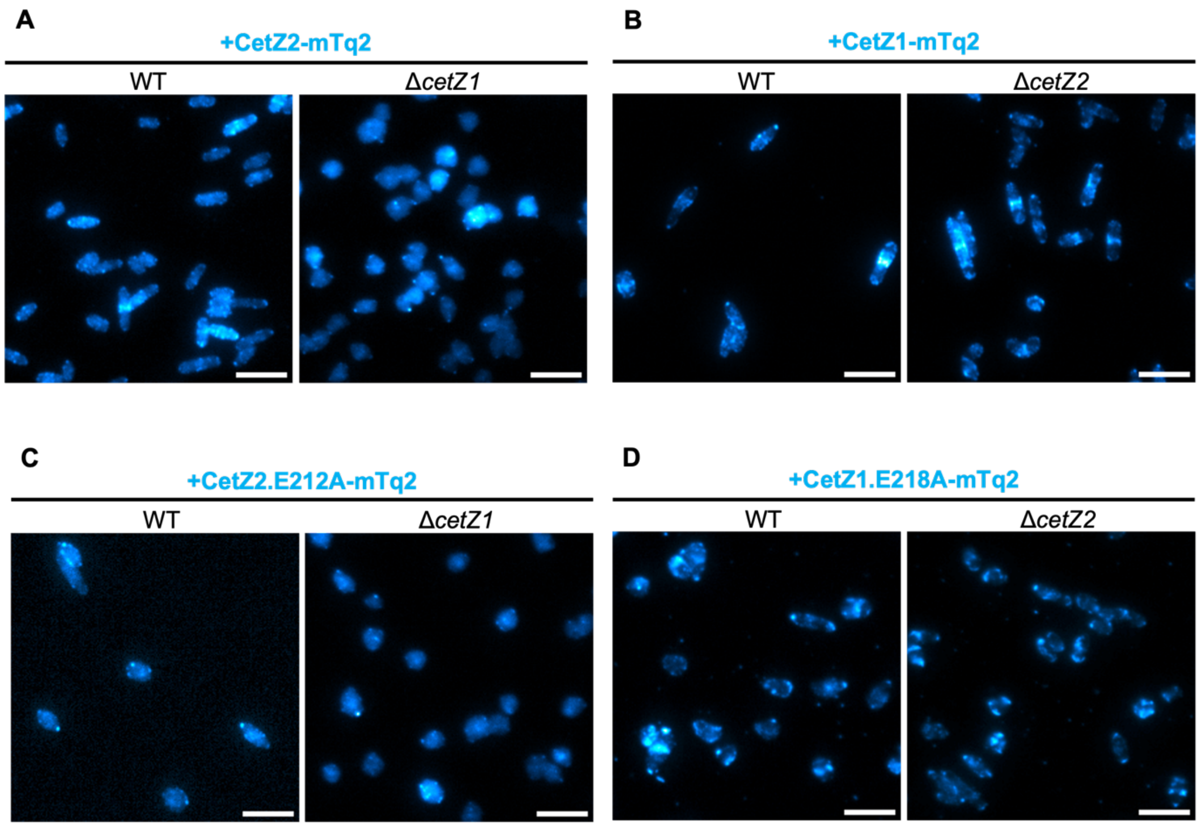


**Supplementary Figure S5.** ***Localisation of CetZ1, CetZ2, CetZ1.E218A, and CetZ2.E212A during mid-log growth.*** All strains were grown in Hv-YPCab medium supplemented with 0.2 mM ʟ-tryptophan and imaged at 24 h. Scale bar=5 µm. **A)** WT and Δ*cetZ1* + pHJB6 (CetZ2-mTq2), **B)** WT and Δ*cetZ2* + pHVID135 (CetZ1-mTq2), **C)** WT and Δ*cetZ1* + pHJB12 (CetZ2.E212A-mTq2), and **D)** WT and Δ*cetZ2* + pHJB13 (CetZ1.E218A-mTq2).

**Table S1.** Strains used in this study

| **Strain** | **Genotype** | **Source** |
|---|---|---|
| ID621 H26 | Δ*pyrE2,* cured of pHV2 | [1] |
| ID622 Δ*cetZ1* | Δ*cetZ1* Δ*pyrE2,* cured of pHV2 | [2] |
| ID357 Δ*cetZ2* | Δ*cetZ2* Δ*pyrE2,* cured of pHV2 | [2] |
| ID643 *cetZ2-mTq2CHR* | *cetZ2-EG-mTq* Δ*pyrE2,* cured of pHV2 | This study |

**Table S2.** Plasmids used in this study

| Plasmid | Description | Source |
|---|---|---|
| pTA131_2204_IF | For in frame deletion of *cetZ1* | [3] |
| pTA131-*cetZ2* | For in frame deletion of *cetZ2* | [4] |
| pTA131-*cetZ2mTq2* | For chromosomal insertion of EG linker and mTurquoise2 at the C-terminus of *cetZ2* | This study (Genscript) |
| pTA962 | Expression vector containing the p.*tna* promotor, *pyrE2* and *hdrB* genetic markers. | [5] |
| pTA962-*cetZ1* | For expression of *cetZ1* under the control of the p.*tna* promotor | [4] |
| pTA962-*cetZ1*.E218A | For expression of *cetZ1*.E218A under the control of the p.*tna* promotor | [4] |
| pTA962-*cetZ2* | For expression of *cetZ2* under the control of the p.*tna* promotor | [4] |
| pTA962-*cetZ2*.E212A | For expression of *cetZ2*.E212A under the control of the p.*tna* promotor | [4] |
| pHVID20 | pTA962 based plasmid containing EG linker and YPet ORF, used to make pHJB5 and pHJB11 | [3] |
| pHVID21 | pTA962 based plasmid containing EG linker and mTurquoise2 ORF, used to make pHJB6 and pHJB12 | [3] |
| pHVID135 | For expression of *cetZ1*-mTq2 (containing G-linker region) under the control of the p.*tna* promotor | [3] |
| pHJB13 | For expression of *cetZ1*.E218A-mTq2 (containing G-linker region) under the control of the p.*tna* promotor | [6] |
| pHJB6 | For expression of *cetZ2*-mTq2 (containing EG-linker region) under the control of the p.*tna* promotor | [2] |
| pHJB12 | For expression of *cetZ2*.E212A-mTq2 (containing EG-linker region) under the control of the p.*tna* promotor | This study |
| pHJB5 | For expression of *cetZ2*-YPet (containing EG-linker region) under the control of the p.*tna* promotor | This study |
| pHJB11 | For expression of *cetZ2*-YPet (containing EG-linker region) under the control of the p.*tna* promotor | This study |
| pHJB16 | For dual expression of *cetZ2*-YPet (containing EG-linker region) and *cetZ1*-mTq2 (containing G-linker region) under the control of the p.*tna* promotor | This study |
| pHJB14 | For dual expression of *cetZ2*-YPet (containing EG-linker region) and *cetZ1*.E218A-mTq2 (containing G-linker region) under the control of the p.*tna* promotor | This study |
| pHJB18 | For dual expression of *cetZ2*.E212A-YPet (containing EG-linker region) and *cetZ1*-mTq2 (containing G-linker region) under the control of the p.*tna* promotor | This study |
| pHJB43 | For dual expression of untagged *cetZ2* and *cetZ1*-mTq2 (containing G-linker region) under the control of the p.*tna* promotor | This study |
| pHJB44 | For dual expression of untagged *cetZ2*.E212A and *cetZ1*-mTq2 (containing G-linker region) under the control of the p.*tna* promotor | This study |
| pHJB45 | For dual expression of *cetZ2*-mTq2 (containing EG-linker region) and untagged *cetZ1* under the control of the p.*tna* promotor | This study |
| pHJB46 | For dual expression of *cetZ2*-mTq2 (containing EG-linker region) and untagged *cetZ1*.E218A under the control of the p.*tna* promotor | This study |

| **Table S3.** Oligonucleotides used in this study | | |
|---|---|---|
| **Name** | **Sequence 5'-3'** | **Description/use** |
| HfxZ3_F | CCCCCGGGAATTCATATGAAAACCGTCCTGATTG GTGTGGGG | Forward primer for amplification of *cetZ2/cetZ2.E212A* ORFs |
| HfxZ3_RnoS | CGCGGATCCTCACAGCAGGTCGTCGAGGTCGTC | Reverse primer for amplification of *cetZ2/cetZ2.E212A* ORFs, excluding the stop codon |
| BglII_CetZ1 | GGCGGCGCTAGCATGAAGCTCGCAATGATCG | Forward primer for amplification of *cetZ1/cetZ1.E212A* ORFs, incorporating BglII at the 5' end |
| NheI_CetZ1 | GGCGGGCGCTAGCATGAAGCTCGCAATGATC | Forward primer for amplification of *cetZ1/cetZ1.E212A* ORFs, incorporating NheI at the 5' end |
| T3 | GCAATTAACCCTCACTAAAGG | Reverse primer for amplification of *cetZ1/cetZ1.E218A* ORFs. |

**Legends for Supplementary Videos:**

SV1. 3D render of CetZ2-mTq2 in the wild-type background, produced from pHJB6, after 96 h of culturing in Hv-YPCab medium supplemented with 0.2 mM L-tryptophan. Scale bar=0.5 μm.

SV2. CetZ2-mTq2 produced in the wild-type background from pHJB6. Cells were grown in Hv-YPCab medium supplemented with 0.2 mM L-tryptophan and sampled in early, mid-, and late stationary phase (72, 96, and 120 h respectively) for time-lapse microscopy. Images were taken at 1 min intervals for 30 min. Scale bar=5 μm.

SV3. Single cell examples of tracking using TrackMate2 to analyse dynamics of CetZ2-mTq2, produced from pHJB6 in the wild-type background, during mid-stationary phase (as in SV2). Individual cells are shown without (left) and with (right) traces (shown in yellow) generated by TrackMate2. Scale bars=5 μm.

SV4. Time-lapse microscopy showing CetZ2.E212A-mTq2 produced in the wildtype background from pHJB12. Cells were grown in Hv-YPCab medium supplemented with 0.2 mM L-tryptophan and sampled in mid-stationary phase (96 h). Images were taken at 1 min intervals for 30 min. Scale bar=5 μm.

SV5. Time-lapse microscopy showing CetZ2-mTq2 produced in the wildtype background (left) and Δ*cetZ1* background (right) from pHJB6. Cells were grown in Hv-YPCab medium supplemented with 0.2 mM L-tryptophan and sampled in mid-stationary phase (96 h). Images were taken at 1 min intervals for 30 min. Scale bar=5 μm.

SV6. Time-lapse microscopy showing CetZ2.E212A-mTq2 produced in the wildtype background (left) and Δ*cetZ1* background (right) from pHJB12. Cells were grown in Hv-YPCab medium supplemented with 0.2 mM L-tryptophan and sampled in mid-stationary phase (96 h). Images were taken at 1 min intervals for 30 min. Scale bar=5 μm.

SV7. Time-lapse microscopy showing CetZ1-mTq2 produced in the wildtype background (left) and Δ*cetZ2* background (right) from pHVID135. Cells were grown in Hv-YPCab medium supplemented with 0.2 mM L-tryptophan and sampled in mid-stationary phase (96 h). Images were taken at 1 min intervals for 30 min. Scale bar=5 μm.

SV8. Time-lapse microscopy showing CetZ1.E218A-mTq2 produced in the wildtype background (left) and Δ*cetZ2* background (right) from pHJB13. Cells were grown in Hv-YPCab medium supplemented with 0.2 mM L-tryptophan and sampled in mid-stationary phase (96 h). Images were taken at 1 min intervals for 30 min. Scale bar=5 μm.

SV9. Time-lapse microscopy showing wildtype cells containing pHJB43 for production of CetZ1-mTq2 and CetZ2 (left) or pHJB44 for the production of CetZ1-mTq2 and CetZ2.E212A (right). Cells were grown in Hv-YPCab medium supplemented with 0.2 mM L-tryptophan and sampled in mid-stationary phase (96 h). Images were taken at 1 min intervals for 30 min. Scale bar=5 µm.

SV10. Time-lapse microscopy showing wildtype cells containing pHJB45 for production of CetZ2-mTq2 and CetZ1 (left) or pHJB46 for the production of CetZ2-mTq2 and CetZ1.E218A (right). Cells were grown in Hv-YPCab medium supplemented with 0.2 mM L-tryptophan and sampled in mid-stationary phase (96 h). Images were taken at 1 min intervals for 30 min. Scale bar=5 µm.
